# Combining statistical and neural network approaches to derive energy functions for completely flexible protein backbone design

**DOI:** 10.1101/673897

**Authors:** Bin Huang, Yang Xu, Haiyan Liu

## Abstract

A designable protein backbone is one for which amino acid sequences that stably fold into it exist. To design such backbones, a general method is much needed for continuous sampling and optimization in the backbone conformational space without specific amino acid sequence information. The energy functions driving such sampling and optimization must faithfully recapitulate the characteristically coupled distributions of multiplexes of local and non-local conformational variables in designable backbones. It is also desired that the energy surfaces are continuous and smooth, with easily computable gradients. We combine statistical and neural network (NN) approaches to derive a model named SCUBA, standing for Side-Chain-Unspecialized-Backbone-Arrangement. In this approach, high-dimensional statistical energy surfaces learned from known protein structures are analytically represented as NNs. SCUBA is composed as a sum of NN terms describing local and non-local conformational energies, each NN term derived by first estimating the statistical energies in the corresponding multi-variable space via neighbor-counting (NC) with adaptive cutoffs, and then training the NN with the NC-estimated energies. To determine the relative weights of different energy terms, SCUBA-driven stochastic dynamics (SD) simulations of natural proteins are considered. As initial computational tests of SCUBA, we apply SD simulated annealing to automatically optimize artificially constructed polypeptide backbones of different fold classes. For a majority of the resulting backbones, structurally matching native backbones can be found with Dali Z-scores above 6 and less than 2 Å displacements of main chain atoms in aligned secondary structures. The results suggest that SCUBA-driven sampling and optimization can be a general tool for protein backbone design with complete conformational flexibility. In addition, the NC-NN approach can be generally applied to develop continuous, noise-filtered multi-variable statistical models from structural data.

Linux executables to setup and run SCUBA SD simulations are publicly available (http://biocomp.ustc.edu.cn/servers/download_scuba.php). Interested readers may contact the authors for source code availability.

## 1. Introduction

In past decades, substantial progresses have been made in computational protein design,(Chen, et al., 2018; Dahiyat and Mayo, 1997; Huang, et al., 2016; Kuhlman, et al., 2003) with automated sequence design tools maturing(Alford, et al., 2017; Dahiyat and Mayo, 1997; Gainza, et al., 2016; Liu and Chen, 2016; Xiong, et al., 2014) and examples of successfully designed proteins with *de novo* backbones increasing. On the other hand, current methods for designing protein backbones still heavily rely on structure type-specific heuristic rules or parametric models.(Grigoryan and Degrado, 2011; Huang, et al., 2014; Jacobs, et al., 2016; Lin, et al., 2015; Yeh, et al., 2018) To take full advantage of the plasticity of protein backbone conformations in protein design,(Huang, et al., 2016; MacDonald and Freemont, 2016) it is highly desirable to have a general method that can be used to sample and optimize designable polypeptide backbones without pre-specified amino acid sequences. While a few previous computational studies have suggested that well-folded protein conformations may correspond to minima on free energy surfaces of backbones modeled without specific sidechain information,(Cossio, et al., 2010; Hoang, et al., 2004; Kukic, et al., 2015; Taylor, et al., 2009; Zhang, et al., 2006) most of these studies have aimed at coarsely contouring the free energy landscapes of polypeptides rather than at obtaining accurate backbone structures to be used for amino acid sequence design. One exception was the study of MacDonald et al., in which they developed a C_α_-atom-based statistical energy function that emphasized on the accurate modeling of local backbone conformation, i.e., the backbone conformations of a few consecutive residues.(MacDonald, et al., 2010) The minima of this energy function determined without specific sidechain information have been shown to resemble experimentally-determined loop structures in native as well as in designed proteins.(MacDonald, et al., 2016; MacDonald, et al., 2013) More recently, we have proposed a sidechain-independent statistical model named tetraBASE to model the through-space packing between backbone positions contained in different rigid secondary structure elements (SSEs).(Chu and Liu, 2018) The minima on the tetraBASE energy surface could reproduce various multi-SSE architectures in native proteins, with atomic positional root mean square deviations (RMSD) mostly between 1.5 to 2.5 Å. The tetraBASE model, however, does not describe the internal flexibility of SSEs or the conformation of loops. It is also discontinuous with respect to conformational changes.

Here we report a comprehensive statistical energy function for completely flexible protein backbone conformation sampling and optimization. The model is named SCUBA, standing for SideChain-Unspecialized-Backbone-Arrangement, as sidechains have been considered mainly as steric space holders in the model, so that protein backbones can be sampled and optimized with generally simplified amino acid sequences. This distinguishes SCUBA from existing statistical potentials developed for the modeling or evaluation of protein structures with specific amino acid sequences.(Dong, et al., 2013; Liu, et al., 2014; Ramon Lopez-Blanco and Chacon, 2019; Sippl, 1990; Xu, et al., 2017; Zhou and Zhou, 2002)

A distinct feature of SCUBA is that each of its statistical energy terms depends on a multiplex of geometric variables. To consider multiplex variables jointly should be important because a range of many-body effects may not be reproduced well by a summation of simple terms that depend on only one or two variables.(Chu and Liu, 2018; Xiong, et al., 2014) In practice, the construction of multi-variable statistical energies are challenging,(Liu, et al., 2014; Ramon Lopez-Blanco and Chacon, 2019; Xu, et al., 2017) being associated with technical difficulties such as how to evaluate properly-gauged statistical energies from training data that are unevenly distributed in non-orthogonal and non-isometric multivariable spaces, how to choose appropriate multi-dimensional functional forms to represent the energy surface, and how to reach at continuous models with easily computable gradients.

In the current work, we introduce a general approach that solve the above difficulties. The method, named NC-NN, comprises using adaptive-cutoff neighbor-counting (NC) to estimate properly gauged high-dimensional statistical energies, followed by representing the high-dimensional statistical energy surfaces as neural networks (NN).(Behler and Parrinello, 2007; Galvelis and Sugita, 2017; Lemke and Peter, 2017; Shen and Yang, 2018) The energy terms obtained by this NC-NN approach have analytical gradients, allowing them to be used directly to drive (stochastic) molecular dynamics simulations. The SCUBA model contains NC-NN-derived energy terms to describe the main chain local conformation, the main chain through-space packing, the backbone-dependent side chain conformation, and so on, the relative weights of different energy components calibrated on the basis of SCUBA-driven stochastic dynamics (SD) simulations of natural proteins. The model is then validated by comparing backbones of native proteins with backbones artificially constructed and automatically optimized using SCUBA.

## 2. Methods

### 2.1 The composition of the SCUBA energy function

The total energy is written as the sum of a sidechain-independent and a sidechain-dependent part, namely,

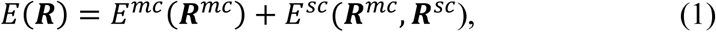

in which ***R***^*mc*^ and ***R***^*sc*^ refer to the atomic coordinates of the main chain atoms and the side chain atoms, respectively.

The sidechain independent part has been defined as the sum of four components,

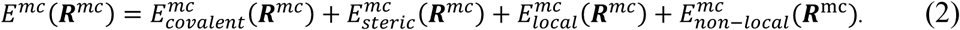

The covalent component 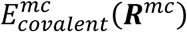 consists of harmonic bond length, bond angle, and improper dihedral angle terms. The steric component 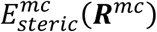 is a sum over main chain atom pair distance dependent terms. The local conformation component 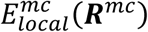 is defined as a sum over windows centered at individual residue positions, namely,

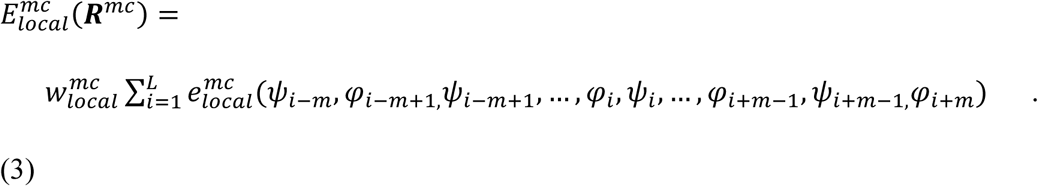

The 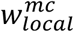 is a weighting factor. For each residue position *i*, the term 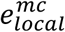 depends on a series of consecutive Ramachandran torsional angles along the peptide chain centered around *i*, namely, from *ψ*_*i*−*m*_ to *φ*_*i+m*_. We use *m=0* for the first and the last two positions of a peptide chain (i.e., *i*≤ 2 or *i*≥ *L* − 1), and *m=2* for middle positions. For the latter positions, the 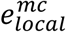 is decomposed into a single residue Ramachandran term and a multi-residue correlation term,

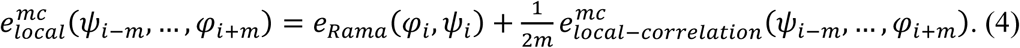

If we only keep the *e*_*Rama*_ (*φ*_*i*_,*ψ*_*i*_) term on the right side of formula (4), we would be ignoring correlations between neighboring backbone positions. Besides the energy terms defined by formulae (3) and (4), an explicit Cartesian coordinate-dependent main chain hydrogen bond term 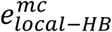 can be optionally added to 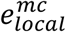 to improve hydrogen bonding geometries in helices (see Supplementary Methods).

The through-space component 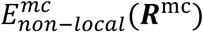 in formula (2) is treated as a sum over residue pairs that are separated by at least 4 residues in the sequence, namely,

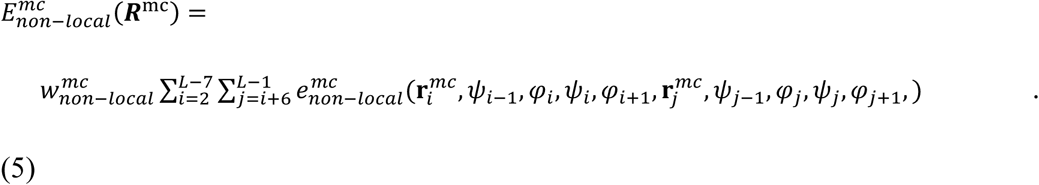

The 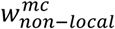 is a weighting factor. In formula (5), the residue pairwise interaction 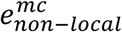 depends not only on the coordinates of all the main chain atoms at positions *i* and *j* (noted as 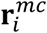 and 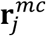, respectively), but also on the local conformations at these two positions as specified by the Ramachandran angles.

The sidechain dependent energy in formula (1) has been defined as the sum of three components,

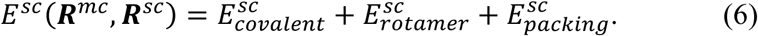

The covalent component 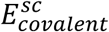 contains usual harmonic terms depending on bond lengths, bond angles and improper dihedral angles. The rotamer component is a sum over residue-wise terms,

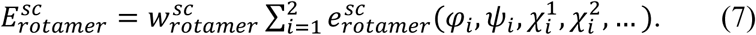

The 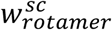 is a weighting factor. For each residue *i*, the rotamer energy depends on not only the sidechain torsional angles 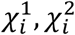, and so on, but also the local backbone conformation as specified by the Ramachandran torsional angles. The component 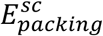 has been treated as a sum over simple distance-dependent atomic pairwise terms (see Supplementary Methods) multiplied by a weighting factor 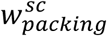. In SCUBA, the main purpose of considering the 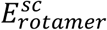 and 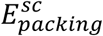 components is to model the steric volume effects of sidechains which may play indispensable roles in shaping the backbone conformational landscape. In addition, the 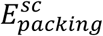 contains inter-atomic attractions (see Supplementary Methods) to counterbalance the thermal expansion effects in finite temperature SD simulations. Other than these, the descriptions of sidechain interactions that are more residue-type-specific, such as the electrostatic interactions, hydrogen-bonding, (de)solvation, and so on, have been intentionally omitted or simplified. The purpose is to minimize the differences between different specific sidechain types, so that the model can be applied to backbones with generic or simplified amino acid sequences.

The statistical energy terms in SCUBA have been derived from more than 10,000 non-redundant training native protein structures (X-ray structure resolution higher than 2.5 Å and sequence identity below 50%).(Wang and Dunbrack, 2005; Xiong, et al., 2014) More details about the SCUBA energy terms are given in Supplementary Methods.

### 2.2 The NC-NN approach to construct high dimensional statistical energies

A general approach has been applied to derive statistical energy terms in SCUBA that each depends on a multiplex of geometric variables, the terms including *e*_*Rama*_, 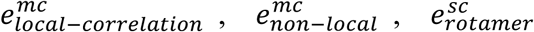, and the optional 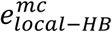 (See Supplementary Methods). This NC-NN approach consists of a neighbor-counting (NC) step followed by neural network-fitting (NN).

#### (1) The neighbor-counting (NC) step

We denote the multi-dimensional geometric variables collectively as ***Θ*** ≡ (*θ*_*i*_, *θ*_2_,…, *θ*_*d*_), and consider two probability density functions in the ***Θ*** space, one denoted as *ρ*^*t*^(***Θ***) corresponding to the distribution of the training data, and the other denoted as *ρ*^*r*^(***Θ***) corresponding to a background or reference distribution. An effective statistical energy as a function of ***Θ*** can be defined as

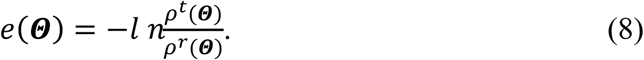

We do not try to determine *e*(***Θ***) through any sort of parametrically estimated *ρ*^*t*^(***Θ***) Instead, for any given point ***Θ***^***x***^, the value of *e*(***Θ***^***x***^) is directly estimated from two set of sample points distributed in the ***Θ*** space. One set denoted as *S*^*t*^ contains samples whose distribution follows *ρ*^*t*^(***Θ***). For a SCUBA energy term, this sample set consists of data extracted from the training native proteins. The other set denoted as *S*^*r*^ consists of samples computationally generated according to the distribution *ρ*^*r*^(***Θ***). Then the ratio 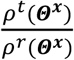 can be estimated as the ratio between the (properly normalized) numbers of neighboring points of ***Θ***^***x***^ in the *S*^*t*^ set and in the *S*^*r*^ set, respectively. Namely,

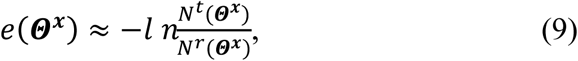

in which *N* ^*t*^ and *N* ^*r*^ represent the respective normalized numbers of neighbors. We note that the different geometric variables constituting ***Θ*** do not need to be orthogonal, to be isometric with respect to the Cartesian coordinates, or to be linearly independent from each other.

For formula (9) to be meaningful, for any given ***Θ***^***x***^ point, the values *N*^*t*^ and *N*^*r*^ should be estimated in exactly the same way. This requirement fulfilled, the ratio form in formula (9) should lead the computed energy to be relatively insensitive to the exact way of how neighbor counting has been implemented. A general way to compute the normalized number of neighbors *N* in a sample set *S* for the probing point ***Θ*** ^***x***^ is to define the following multi-dimensional kernel function

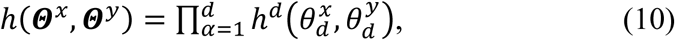

in which the one dimensional function 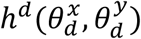 takes the value of 1 if the difference between 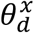 and 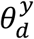 is below a (adaptively) chosen cutoff, and gradually goes to zero as the difference increases (the “soft” cutoff approach). Given *h*, the normalized number of neighbors *N* of point ***Θ*** ^***x***^ in set *S* can be computed as

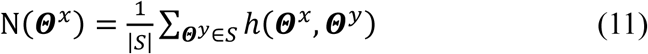

in which |*S*| stands for the cardinal or the number of points in *S*. As the distribution of training points in the ***Θ*** space is usually extremely uneven, the kernels 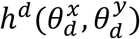 may need to be adaptively defined, associated with larger cutoffs in sparsely populated ***Θ*** regions to reduce statistical uncertainties, and with smaller cutoffs in densely populated ***Θ*** regions to increase resolution.(Xiong, et al., 2014)

#### (2) The neural-network (NN) fitting step

A major drawback of the above neighbor-counting (NC) approach is that it is computationally too expensive for on-the-fly energy evaluations in conformation sampling or optimization. In addition, the directly estimated energy surfaces are roughed by statistical noises, analytical derivatives of the energy being unavailable.

These drawbacks are overcome after the NN step. Inspired by the idea of replacing first principle potential energy surfaces with artificial neural networks (NNs),(Behler and Parrinello, 2007; Shen and Yang, 2018) we use NNs to represent the NC-derived statistical energies as analytical functions of multiplexes of geometrical variables.(Galvelis and Sugita, 2017; Lemke and Peter, 2017) Here, the inputs of a NN are the geometric variables (i.e. the different constituents of ***Θ***) encoded with chosen encoding schemes. The output is the value of the statistical energy. The NN model is trained using the NC-estimated single point statistical energies at a diversely distributed set of points in the ***Θ*** space. As the training of the NN needs to be carried out only once, the NC estimation can be carried out for as many ***Θ*** space points as needed to provide a sufficient amount of training data to the NN.

The NNs used in the current work are of three-layers, implementing the following mapping from an input encoding vector ***x*** to an output real value,

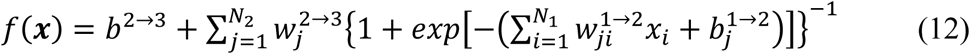

in which *N*_1_ is the number of nodes in the first or input layer, *N*_2_ the number of nodes in the second layer, 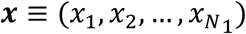 the input vector encoding a point in the ***Θ*** space, the coefficients 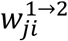 and 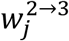 the weights connecting the first and the second layers and the second and the third layers, respectively, and 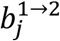 and *b*^2→3^ the respective biases. The input to node *j* in the second layer is 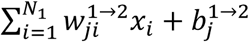, which is mapped to the output by the transformation 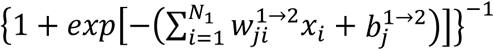. The single node in the third or output layer accepts inputs from all the second layer nodes, combines them linearly, and adds a biasing value to generate the final output.

### 2.3 Calibrating and testing SCUBA by stochastics dynamics simulations of native proteins

The weighting factors in the SCUBA model, including 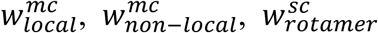, and 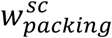, have been introduced to compensate for potentially redundant or double-counted interactions between different energy components. These weights have been calibrated using SD simulations of 33 native proteins, using in-house developed codes implementing SCUBA-driven SD with bond lengths constrained by SHAKE.(Vangunsteren, et al., 1981) The codes can be applied to optimize backbone conformations through simulated annealing, or to test if any conformational minimum on the SCUBA energy surface is stable against thermal fluctuations at a given “temperature” (we assume the statistical energies defined according to formula (8) to be of the physical unit of *k*_*B*_*T*_*0*_, in which *k*_*B*_ is the Boltzmann constant and the temperature *T*_*0*_ is 300 *K*. In later discussions, we will use the reduced temperature *T*_*r*_, with *T*_*r*_ = 1 corresponding to 300 *K*). The energy weight calibrations have been carried out using an approach that is conceptually similar to force field parameter refinements using thermodynamics cycles in conformational space(Cao and Liu, 2008) or “contract divergence”(Jumper, et al., 2017; Várnai, et al., 2013), with the objective of parameterization being to stabilize the native conformational states relative to conformations further away from the native structures. After determining the weights, SD simulations have been carried out on the native proteins in their original sequences as well as in simplified sequences, in which all residues in helices have been changed into leucine, residues in strands changed into valine, and residues in loops removed of sidechains. More details are given in Supplementary Methods.

### 2.4 Designing artificial backbones using SCUBA by SD and simulated annealing

#### (1) Building the initial backbone structures

To design a backbone, an intended “framework” is specified first. This framework defines at a very coarse level the intended backbone architecture, including the numbers, types, sequential orders, and approximate lengths of secondary structure elements (SSE). It also specifies how the SSEs should be organized in the three-dimensional space to follow an abstractive multi-layered form as summarized from native protein folds by Taylor et al.(Taylor, 2002) In an initial structure artificially constructed according to this form, the N or C-terminal end positions of SSEs in the same SSE layer fall on grid points on a line in a 2-dimensional plane. The end-to-end directions of the SSEs are perpendicular to the plane. SSEs in different layers have their ends on different parallel lines on the plane. Given the intended framework, helix and strand fragments of expected lengths have been constructed with the given end positions, with backbone torsional angles randomly drawn from distributions associated with respective SS types, and with given end-to-end directions. Then loops of given lengths have been built using the kinematic closure algorithm(Coutsias, et al., 2004) to link the SSEs in a given order. For a given framework specification, different initial backbones have been built by using different random seeds. A two-stage SD simulated annealing procedure (see Supplementary Methods) has been applied to optimize each artificially constructed backbone. In the first stage, no sidechains have been considered. In the second stage, leucine (valine) sidechains have been considered for all helix (strand) positions. For each intended framework, 10 conforming final SCUBA-optimized structures have been obtained. Each final structure has been used as a query to search against the protein data bank (PDB) using the Dali server(Holm and Laakso, 2016) (or the mTM-align server(Dong, et al., 2018) if the Dali search did not return any matching structure with a Z-score above 6.0)

## Results and discussions

### 3.1 Statistical energy terms constructed by the NC-NN approach

Despite that the construction of a multi-dimensional NC-NN term involves many steps with intricate details (see Supplementary Methods), the NC-NN approach seems to be robust: by simply following common senses and using not excessively fine-tuned parameters for the intermediate steps, a final statistical energy term may be obtained to faithfully model a high-dimensional native distribution of a multiplex of strongly correlated geometric variables. This point is visually illustrated in Supplementary Figures S1 to S7, which show several examples of distributions of the NC-estimated statistical energies (Figures S1 to S3) and comparisons between the NC-estimated and the NN-estimated energies (Figures S4 to S7). Brief discussions of the implications of the presented data have been included in corresponding figure captions.

### 3.2 Calibrating the energy function by SD simulations of native proteins

In Figure 1, the averaged RMSDs of the structures sampled in the SCUBA-driven SD simulations from respective native structures are given. The results from four sets of simulations have been compared. In the first three sets of simulations, the test proteins have their native sidechains. The medium value of the averaged RMSDs for the 33 simulated proteins is 1.85 Å when both the optional main chain local hydrogen bond terms and the radius of gyration restraint are turned off (see Supplementary Methods). The medium RMSD is reduced slightly to 1.6 Å upon the inclusion of the optional main chain local hydrogen bond terms, and further reduced to 1.25 Å upon the additional inclusion of the radius of gyration restraint. There does not seem to be any systematic difference between the RMSDs obtained for backbones of different fold classes. The last set of simulations have been carried out on the proteins of simplified sequences. The medium RMSD is 2.23 Å when both the main chain local hydrogen bond terms and the radius of gyration restraint are turned on. If these optional potentials are turned off, the medium RMSD is slightly larger (2.47 Å). These simulations have been carried out with the finalized set of energy weights given in Supplementary Methods. The effects of varying the individual or the overall energy weights on the RMSDs have been summarized in supplementary Figures S8 and S9, with brief discussions of the implications of the presented data included in corresponding figure captions.

**Figure 1.**
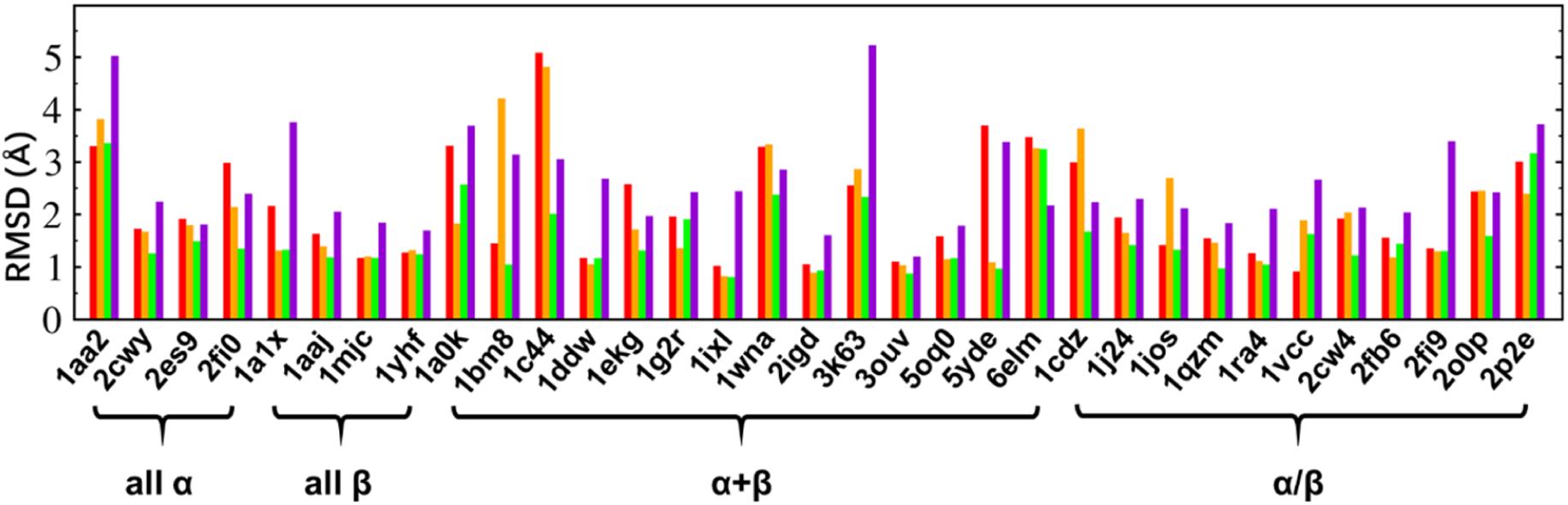
The averaged RMSDs from native structures of structures sampled in the SCUBA-driven SD simulations using the calibrated energy weights. For each native structure with the given PDB ID, results of four simulations are plotted in different colors. Three simulations have been carried out on the native sequence, with both the main chain local hydrogen bond terms 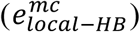 and the radius of gyration (***R***_*g*_) restraint off (red), or 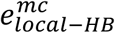 on and the ***R***_*g*_ restraint off (orange), or both 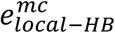 and the ***R***_*g*_ restraint turned on (green). One set of simulations have been carried out on the simplified sequence, with both 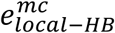 and the ***R***_*g*_ restraint turned on (violet). The fold classes are indicated below the PDB IDs.

The above results suggest that with the chosen set of energy weights, most of the native protein backbones are stable against thermal fluctuations at *T*_*r*_ = 1.***Θ***, and the SD sampled conformations remain close to the native structures. In addition, when the native sequences have been substituted with the generally simplified sequences, the stability of most native backbones can still be retained, with the RMSD medium values still below 2.5 Å.

### 3.3 Backbones optimized from artificial initial structures and their similarity to native structures

The intended frameworks for artificial backbone construction are given in Tables 1 to 4 with framework IDs and intended types, orders, and lengths of the secondary structure elements (SSEs). These frameworks cover different types of SSEs interacting with each other in various combinations and relative geometries. Examples of initial and SCUBA-optimized structures for each framework are given in Figures 2 to 4. The framework in Table 1 is a 3-helix bundle. Frameworks in Table 2 comprise four SS segments, including a four-helix bundle, a 4-strand β sheet, and several two layered frameworks containing one helix packed against a 3-strand β sheet. Frameworks in Table 3 comprise six SS segments arranged in two layers, each containing a two-helix layer packed against a 4-strand β sheet. Frameworks in Table 4 comprise six SS segments arranged into three layers, each containing two single-helix layers packed at the two opposite sides of a 4-strand β sheet.

**Table 1.**
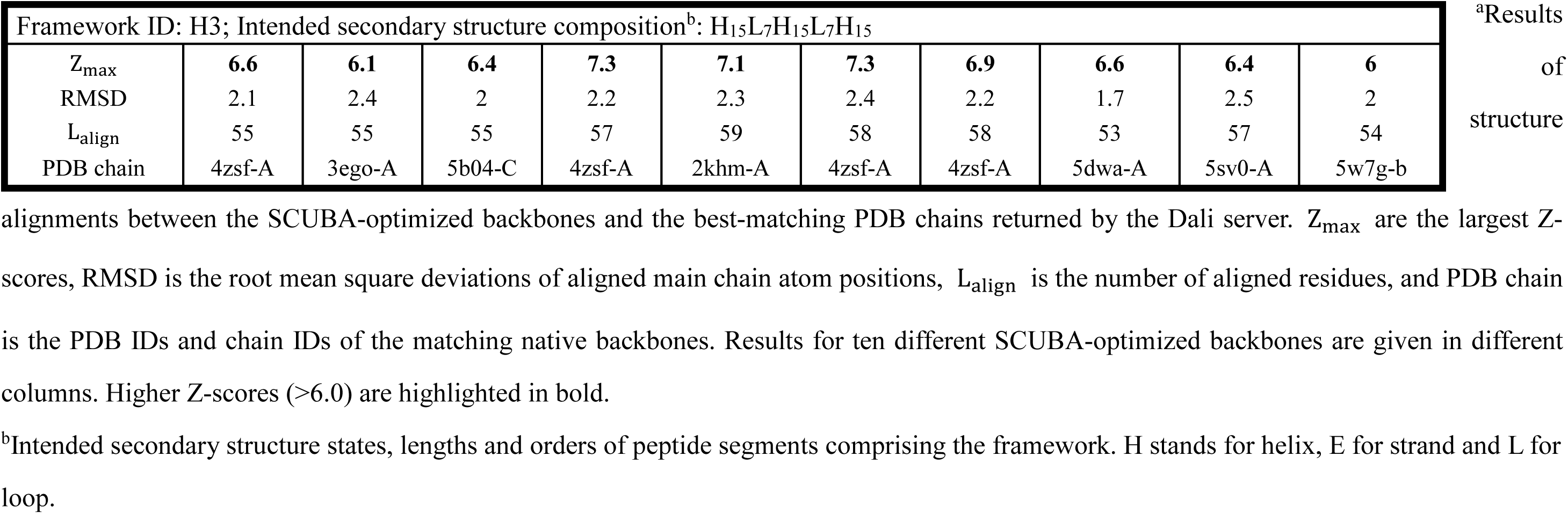
PDB search results using the SCUBA-optimized backbones obtained for a Framework consisting of three helices.^a^

**Table 2.**
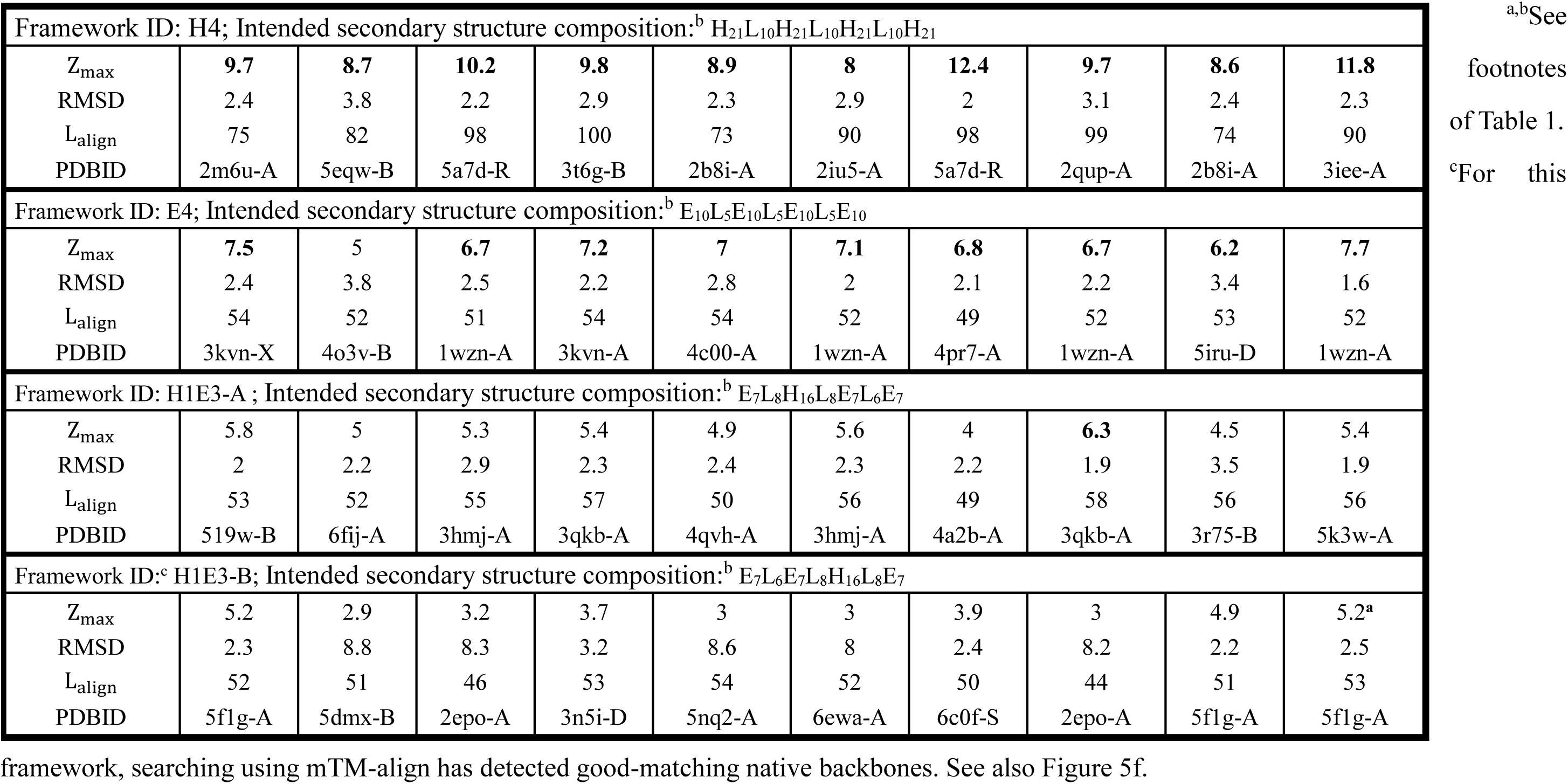
PDB search results using SCUBA-optimized backbones obtained for Frameworks consisting of four secondary structure segments.^a^

**Table 3.**
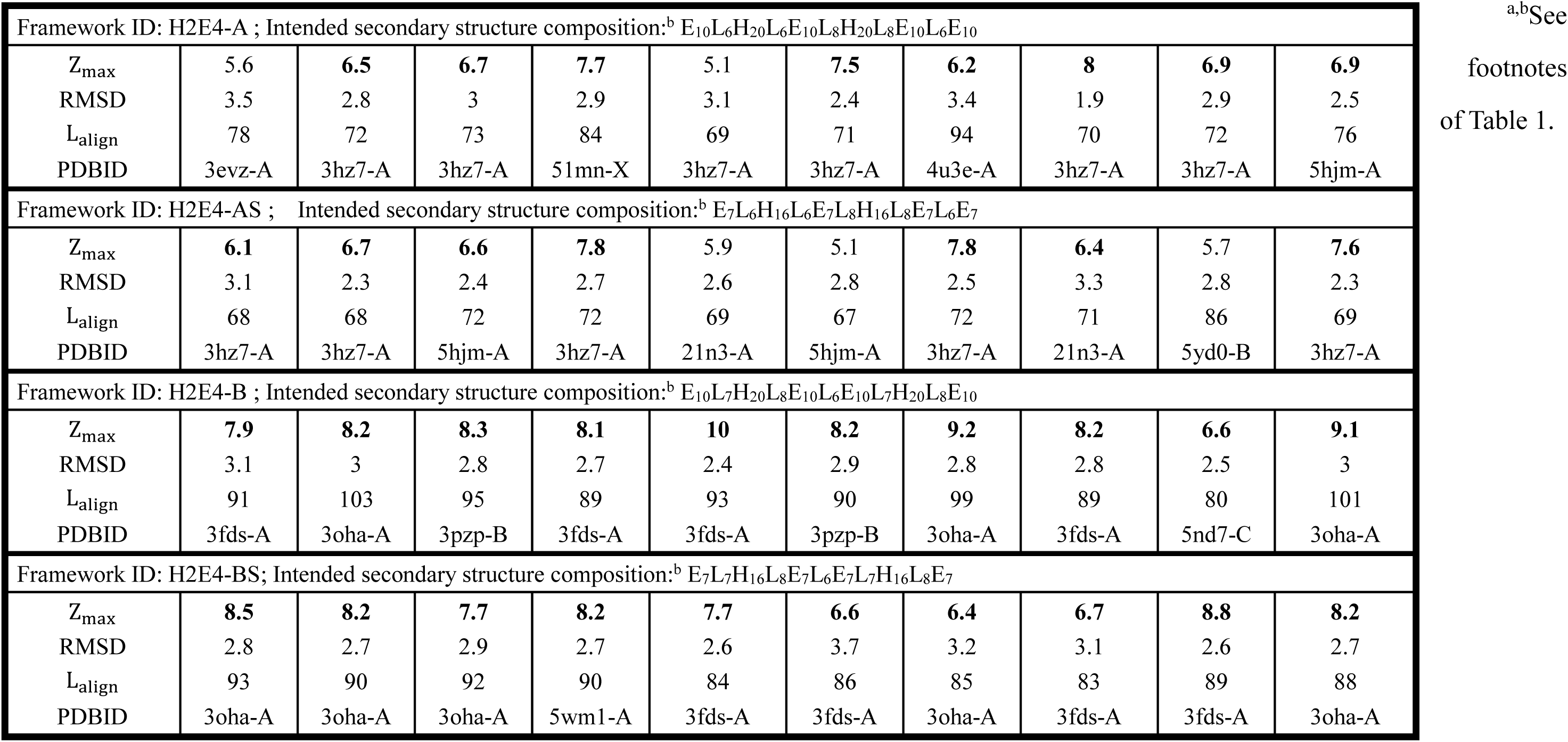
PDB search results using SCUBA-optimized backbones obtained for Frameworks consisting of six secondary structure segments arranged in two layers.^a^

**Table 4.**
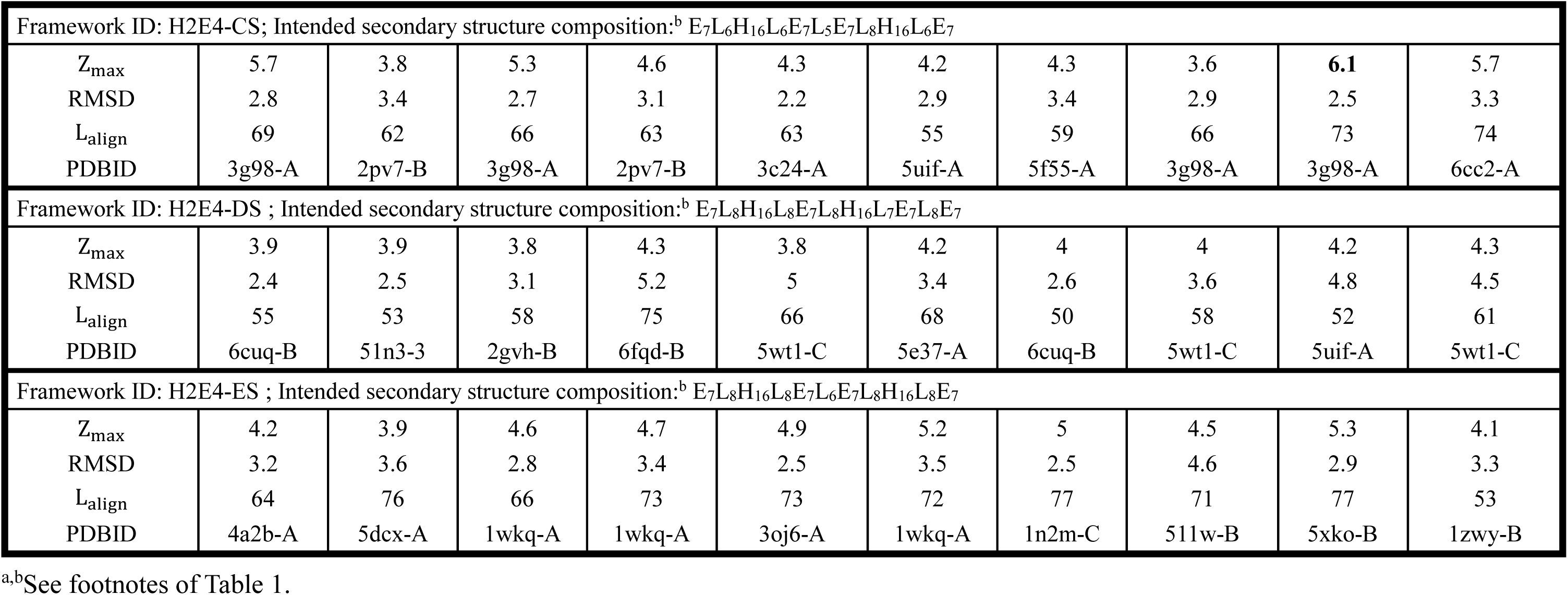
PDB search results using SCUBA-optimized backbones obtained for Frameworks consisting of six secondary structure segments arranged in three layers.^a^

**Figure 2.**
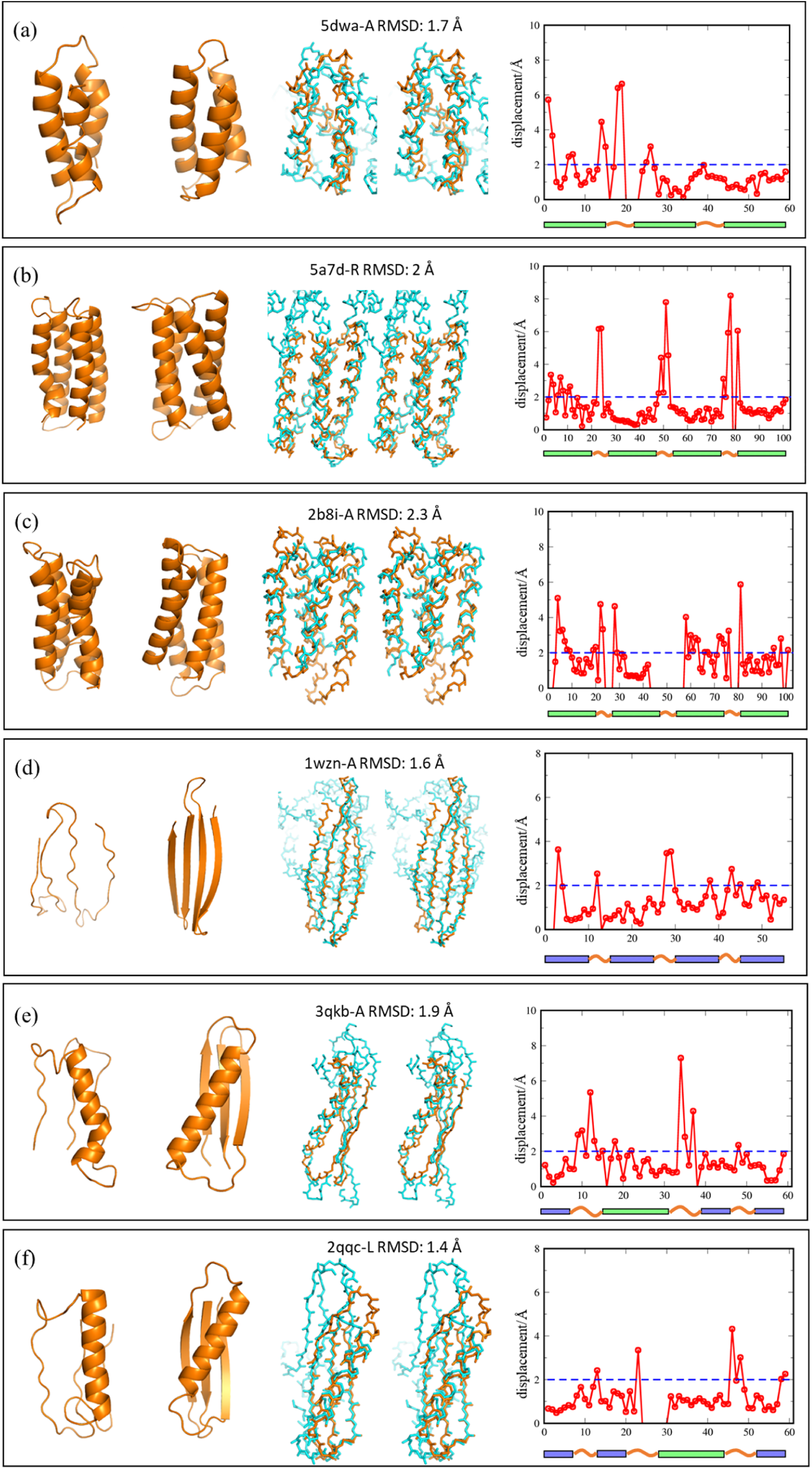
From left to right in each panel, artificial initial structure, SCUBA-optimized structure, stereo view of SCUBA-optimized structures superimposed with a matching native PDB structure, and displacements of aligned positions between the SCUBA-optimized and the matching native PDB backbone. Artificial structures are shown in orange. Native structure are shown in light blue. PDB IDs, chain IDs and overall RMSDs are given above the stereo views. Secondary structure segments are indicated under the displacement plots (Helix: green box. Strand: blue box. Coil: red line.). (a) Framework H3. (b) Framework H4, SCUBA-optimized structure in left-handed twist. (c) Framework H4, SCUBA-optimized structure in right-handed twist. (d) Framework E4. (e) Framework H1E3-A. (f) Framework H1E3-B.

In Tables 1 to 4, the PDB IDs of the best-matching native structures (according to the Dali Z-scores) of the SCUBA-optimized backbones are given, together with lengths, Z-scores, and RMSDs of the respective structure alignments. Examples of aligned SCUBA-optimized backbones and native backbones are shown in Figures 2 to 4 in stereo views, together with plots of the displacements of individual aligned positions. The shown native backbones have been found by Dali, except for the one shown in Figure 2f, which has been found by mTM-align.

For the majority of the frameworks in Tables 1 to 3, the SCUBA-optimized backbones can be aligned to one or more native backbones with Dali Z-scores > 6.0. For these high Z-score alignments, the overall RMSDs of the aligned positions are mostly between 2 to 3 Å, with the main chain atom displacements for residue positions contained in regular SSEs being mostly below 2 Å (see the displacement plots in Figures 2 and 3). Especially, for the framework H4, the SCUBA-optimized backbones exhibit inter-helix twist of two types of handedness. Both the backbone exhibiting left-handed twist (Figure 2b) and the one exhibiting right-handed twist (Figure 2c) can be well aligned with known native backbones. The SCUBA-optimized backbone of the 4-strand antiparallel β sheet shown in Figure 2d accurately reproduces the intra-strand twisting and the surface curving of a native β-sheet.

**Figure 3.**
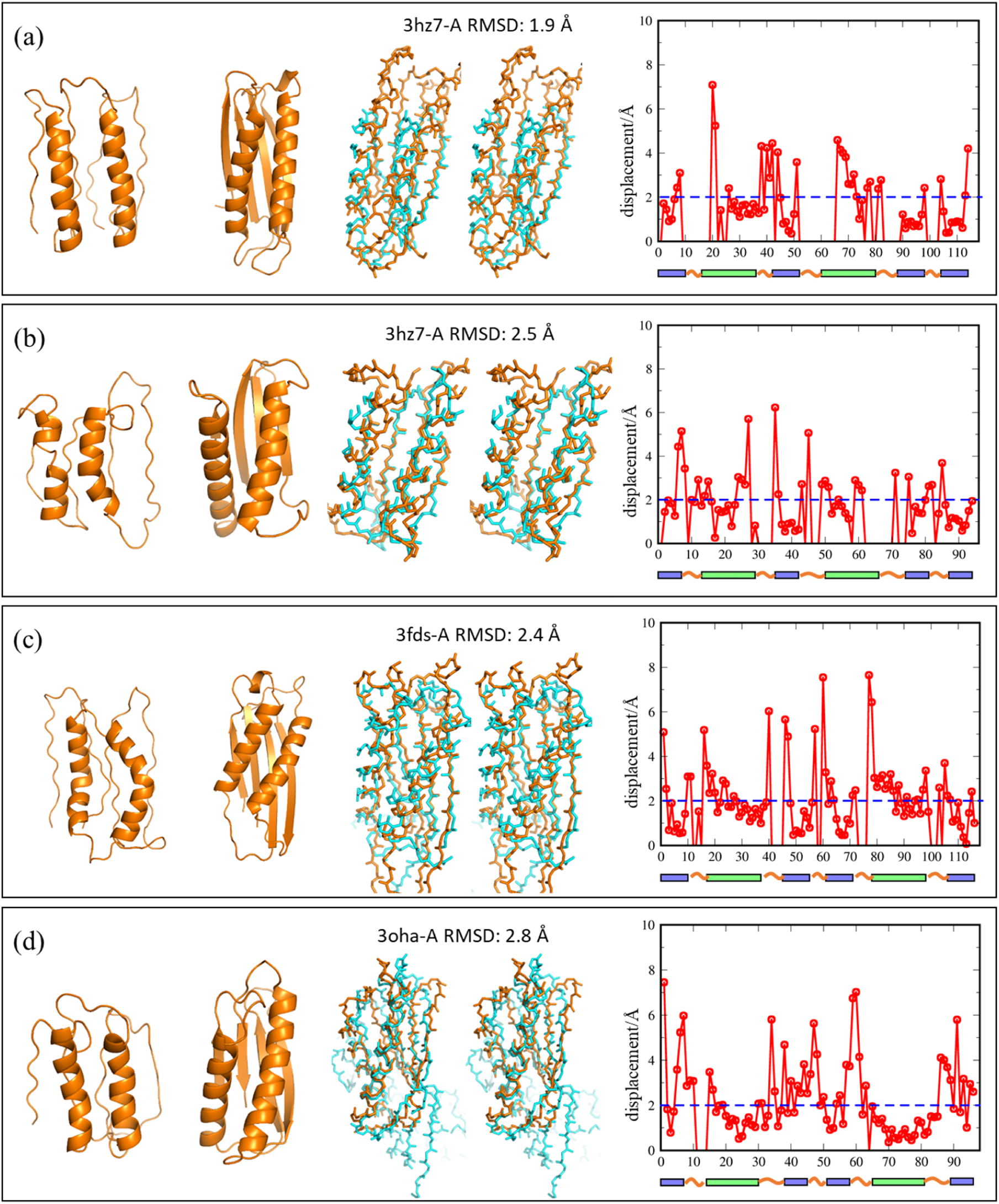
As Figure 2, but for frameworks defined in Table 3. (a) Framework H2E4-A. (b) Framework H2E4-AS. (c) Framework H2E4-B. (d) Framework H2E4-BS.

For the frameworks in Table 4, matching native structures with Dali Z-scores above 6.0 have been detected for only one SCUBA-optimized structure obtained for the framework H2E4-CS (Figure 4a). For the other SCUBA-optimized backbones in Table 4, similar native backbones with high Dali Z-scores have not been found, even though the SCUBA-optimized backbones obtained for these frameworks seem to have similarly well-formed and packed SSEs (Figures 4b and 4c) as those obtained for the other frameworks. It could be possible that the particular SSE arrangements specified for these frameworks were of relatively poor designability.

**Figure 4.**
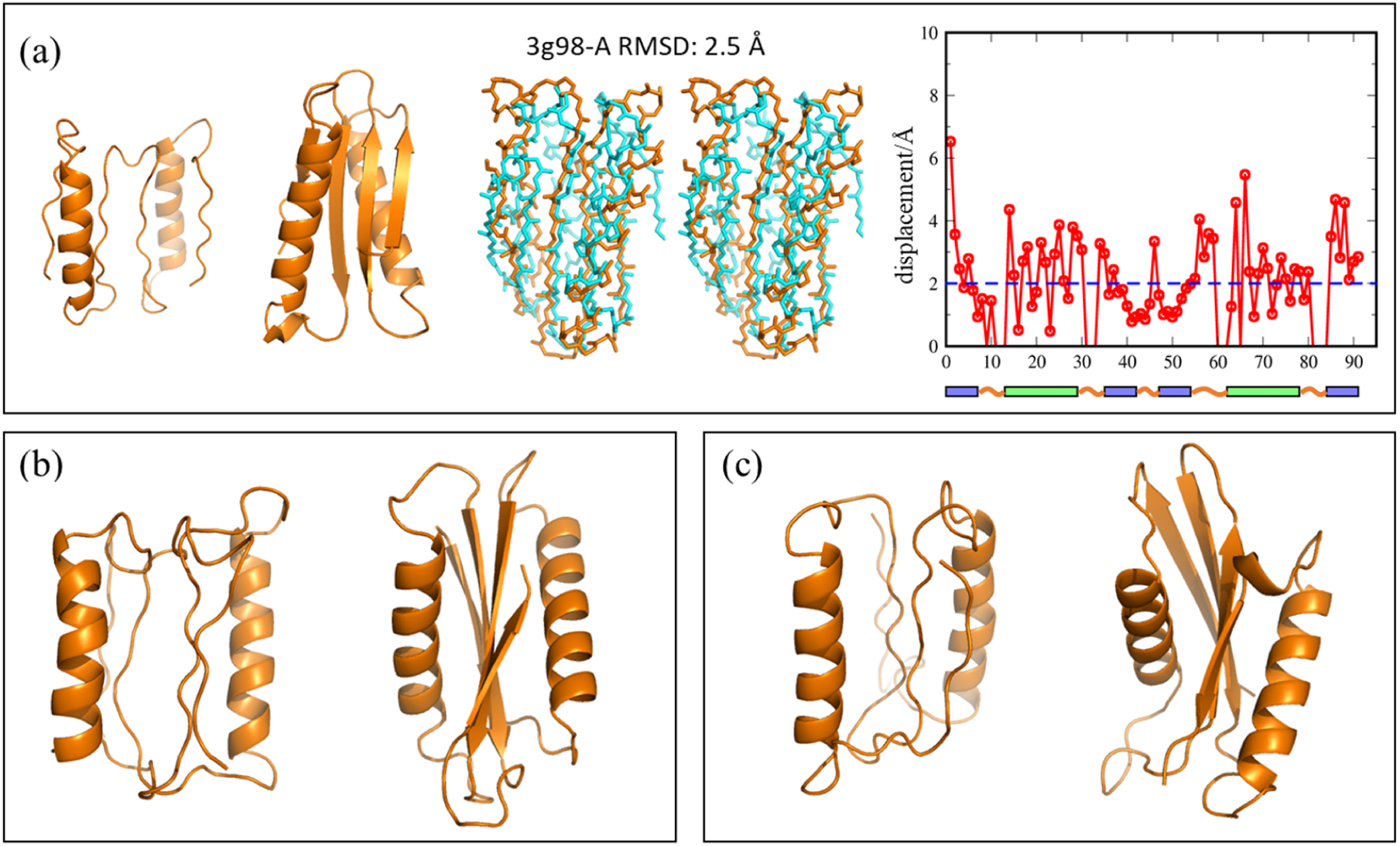
(a) As Figure 2, but for framework H2E4-CS. (b) An initial structure and a SCUBA-optimized structure for framework H2E4-DS. (c) As (b) but for framework H2E4-ES.

## 4. Conclusions

SCUBA as a statistically-learned model of protein conformations is distinct from existing ones in both derivations and outcomes. It can be applied to sample and optimize protein backbones with complete conformational flexibility. With sidechains mainly serving as space holders in SCUBA, generic amino acid sequences may be used in place of specific ones. The good agreements between the artificially-constructed SCUBA-optimized backbones and native backbones suggest that the SCUBA SD approach may potentially be applied to facilitate a variety of protein design tasks, such as to construct *de novo* scaffolds to host functional centers, to restructure backbone segments to form a new active site, or to construct the backbone of a peptide ligand docked onto a receptor. In our future work, we will continue to refine this approach as a protein backbone design tool and to work with collaborators to experimentally verify the designability of the SCUBA-optimized backbones.

SCUBA relies on its various NC-NN-derived energy terms to faithfully reproduce the coupled distributions of multiplexes of conformational variables in designable backbones. The NC-NN approach overcomes common technical difficulties in statistical modeling of highly unevenly distributed data in non-orthogonal and non-isometric multivariable spaces, with results usable in efficient gradient-requiring sampling/optimization algorithms. Although to derive an energy term by NC-NN unavoidably involves some heuristic choices of parameters, the approach is a robust one with the results being relatively insensitive to the exact choices of parameters or hyper parameters. As a general approach to derive multidimensional statistical models from a large amount of structural data, the NC-NN method may be applied to other structural bioinformatics problems besides protein backbone design.

## Acknowledgements

We thank Dr. Bin Jiang for discussions about the neural network potential. This work was supported by the National Natural Science Foundation of China (Grants 21773220 and 31570719).

