## Supplementary Materials for "Combining statistical and neural network approaches to derive energy functions for completely flexible protein backbone design"

**Table of Contents**

1. Supplementary Methods

   1.1 The SCUBA energy terms

   1.2 Calibrating the energy weights by SD simulations

   1.3 Optimizing the artificially constructed backbones by SD simulated annealing

2. Supplementary Table

3. Supplementary Figures

### 1. Supplementary Methods

#### 1. 1. The SCUBA energy terms

**(1) The $e_{Rama}(\varphi,\psi)$ term**

The probability density of the Ramachandran torsional angles of residues in the coil regions of the training native protein structures, $\rho_{\text{rama}}(\varphi, \psi)$, was estimated by dividing the $\varphi$-$\psi$ plane evenly into $5^{o} \times 5^{o}$ areas and computing the frequency of the torsional angles in each area. The energy values at grid points or centers of the areas were estimated as $-ln\frac{\rho_{rama}(\varphi,\psi)}{\rho^{ref}(\varphi,\psi)}$, in which $\rho^{ref}(\varphi, \psi)$ was a uniform distribution. The resulting set of $(\varphi, \psi, e_{rama})$ values have been used to train a neural work $NN_{rama}$ that defines the $e_{Rama}(\varphi, \psi)$ in formula (4) of the main text analytically. As input to $NN_{rama}$, each torsional variable $\tau$ has been encoded by a 6-component vector,

$$\tau \rightarrow (sin\, k\tau\, , cos\, k\tau\, , k = 1,2,4), \tag{S1}$$

which leads to a total number of 12 input components encoding both torsional angles, or $N_{input} = 12$. It is easy to see that the maximum value of $k$ (here 4) determines the finest torsional angle resolution of the resulting energy term. Thus a controlled extent of smoothness has been enforced on the energy function by the encoding scheme to produce the neural network input. The number of nodes in the middle layer, $\text{N}_{\text{middle}}$, is 16. For this particular case concerning $e_{Rama}(\varphi, \psi)$, the data used to train the neural network covers the entire $\varphi$-$\psi$ space thoroughly and the smoothness of the resulting energy surface is enforced by the form of the encoding function. There has been little room left for overfitting, leaving the exact value of $N_{middle}$ not important as long as it is large enough to ensure optimum fitting between the training and trained energy values. The RMS error of fitting by the final $NN_{rama}$ is 0.18, and cannot be reduced by further increasing of $N_{middle}$. In Figure S5, contour maps of the statistically estimated and the analytical $e_{Rama}$ values are compared. This comparison indicates that a large portion of the RMS fitting error has come from noises of the statistically estimated values in $\varphi$-$\psi$ regions associated with higher energies.

**(2) The $e^{mc}_{local-correlation}$ term**

To estimate $\frac{\rho^t}{\rho^r}$ by neighbor counting (NC), the training set $S^t$ consists of points in the eight-dimensional space of $(\psi_{i-2}, \dots, \varphi_{i+2})$ extracted from all pentapeptide fragments in the training native protein structures, while the reference set $S^r$ consists of points computationally sampled according to the reference distribution $\rho^r$, which is the multiplication of the distributions of the Ramachandran angles of individual residues,

$$\rho^r(\psi_{i-2}, \dots, \varphi_{i+2}) = \rho_\psi(\psi_{i-2})\ \left[\prod_{l=-1}^{1} \rho_{rama}(\varphi_{i+l}, \psi_{i+l})\ \right]\rho_\varphi(\varphi_{i+2}), \qquad \text{(S2)}$$

in which $\rho_\psi$ and $\rho_\varphi$ are the respective 1-d marginal distribution densities of the 2-d density $\rho_{rama}$.

For each torsional angle component $\tau$, the 1-d kernel function (corresponding to $h^d(\theta_d^x, \theta_d^y)$ in formula (12) of the main text) used to determine the number of neighbors is of the form

$$\mathrm{h}(\tau^x, \tau^y) = \begin{cases} 1, & if\ \ |\tau^x - \tau^y| < \delta_{min},\ and \\ \left[\frac{(\tau^x - \tau^y)^2 - \delta_{max}^2}{\delta_{min}^2 - \delta_{max}^2}\right]^2, & if\ \ \delta_{min} \le |\tau^x - \tau^y| < \delta_{max},\ and \\ 0, & if\ \ |\tau^x - \tau^y| < \delta_{max}. \end{cases} \qquad \text{(S3)}$$

Here $\tau^x$ is the torsional angle value of the probing point (the point at which the energy is to be evaluated) and $\tau^y$ the torsional angle value of a training point (a point extracted from the training native protein structure). Formula (S3) defines a soft cutoff scheme with cutoff parameters $\delta_{min}$ and $\delta_{max}$ to determine if $\tau^y$ should be counted as a neighbor of $\tau^x$. As discussed in the main text, the cutoffs have been chosen adaptively so that the resulting energy term can be of higher resolution in regions densely populated by training points while of smaller statistical uncertainties in the sparsely populated regions. Specifically, we chose five groups of values for ($\delta_{min}$, $\delta_{max}$), which are ($5^o, 15^o$), ($8^o, 20^o$), ($10^o, 30^o$),($15^o, 45^o$), and ($30^o, 60^o$) in increasing values. For a given probing point, either the smallest cutoffs that lead to an accumulated number of neighbors ($N^t * |S|$ by the notation in the main text) larger than 6.0, or the largest cutoffs, have been used. A small value ($10^{-8}$) were added to the computed $N^t$ and $N^r$ before their ratio was taken to regularize the computed energy in case no neighboring training points could be found with even the largest cutoffs.

The probability densities $\rho_{\psi}$, $\rho_{rama}$, and $\rho_{\varphi}$ in the definition of the reference distribution $\rho^r$ have been estimated using residues in coil regions of the training native proteins. By using this choice without additional measures to treat pentapeptide fragments contained in regular secondary structure elements (SSEs), $e^{mc}_{local-correlation}$ could have significantly overestimated local interactions in regular SSEs, because the apparent local correlations in regular SSEs in globular proteins can be attributed to non-local stabilization interactions. For examples, β-strands are stabilized by inter-strand hydrogen bonds, and α-helices by inter-helix packing and (de) solvation. Estimating $\rho^t$ using the raw training point sets $S^t$ would lead to significant overestimations of local conformational preferences for helices and strands. To offset this effect in the NC-estimated $e^{mc}_{local-correlation}$, the native training pentapeptide fragments have been assigned to different local conformational types (called "protein blocks") based on the eight torsional angles ($\psi_{i-2}, \ldots, \varphi_{i+2}$) according to the classification by de Brevern and coworkers (Offmann, et al., 2007). The fragments assigned to the protein block type “m” are considered as belonging to the center, non-captioning regions of helices (helix state for short) and those assigned to the protein block type “d” considered as belonging to the center, non-captioning regions of strands (strand state for short). When the points in $S^t$ are counted for their contributions to $N^t$, each of those points corresponding to the helix state fragments has been assigned a weight $w_\alpha$ and each of those corresponding to the strand state fragments has been assigned a weight $w_\beta$. With the values of $w_\alpha$ and $w_\beta$ smaller than 1, the computed contributions to the stability of regular SSEs due to local correlations have been lowered. We note that for conformations belonging to the coil state or the helix state, changing $w_\alpha$ or $w_\beta$ has negligible effects on the relative energies between conformations of the same SS state (Figure S10). For local conformations belonging to the strand state the relative ranking between different configurations are also largely reserved (Figure S10). Thus the main effects of varying $\mathrm{w}_\alpha$ or $\mathrm{w}_\beta$ are on the cross-SS state relative energies. Because of the limited effects of these weights and also because of the lack of more objective criteria to fix them to specific values, $\mathrm{w}_\alpha$ or $\mathrm{w}_\beta$ have been treated as adjustable parameters. Models of $e^{mc}_{local-correlation}$ have been constructed with different values of $w_\alpha$ or $w_\beta$ to favor the different types of SSEs to different extents. For the test results reported in this work, the chosen weights are $w_\alpha = 0.1$ and $w_\beta = 0.01$. These choices make the $e^{mc}_{local-correlation}$ of common helix fragments to be comparable to the lowest $e^{mc}_{local-correlation}$ of coil fragments, and the $e^{mc}_{local-correlation}$ of strand fragments to be around zero (see Figure S1a). In other words, we have adopted the hypothesis that in the absence of non-local interactions, the helix conformations are as stable as the most stable coil conformations and the strand conformations are neither favored nor disfavored by local correlations.

The NC-estimated $e^{mc}_{local-correlation}$ values have been used to train the neural network $NN^{mc}_{local-correlation}$, which represents the dependence of $e^{mc}_{local-correlation}$ on the 8 torsional angle variables in an analytical form. Each torsional angle has been encoded into 8 values using the trigonometric functions as in formula (S1), with $k = 1,2,4$ and 8. This leads to 64 input nodes. The number of nodes in the second layer, $N_{middle}$, is chosen to be 72 by trials and errors. The training data for $NN^{mc}_{local-correlation}$ comprises 180000 points taken from $S^t$ and an equal number of points taken from $S^r$. The resulting NN has been tested using 4500 points from the two sets, the test points not used for NN training.

**(3) The $e^{mc}_{non-local}$ term**

The training set $S^t$ for the estimation of $e^{mc}_{non-local}$ by NC consists of main chain residue pairs with inter-$C_\alpha$ distances below 9.5 Å in the training native protein structures. Residue pairs belonging to the same helix or strand or with sequence separations below 6 (i.e., by less than 5 residues) have been excluded. To generate points in the $S^r$ set, one residue from a randomly chosen native pair was randomly translated (within the 9.5 Å cutoff) and rotated. Geometries that lead to steric clashes (clash cutoffs are 2.5 Å for the N-O distance and 3 Å for the other interatomic distances) have been excluded from $S^r$.

In neighbor counting, the intra-position local conformation subspaces and the inter-position relative geometry subspace are considered jointly. To represent the former subspaces, the 4 backbone torsional angles centered at each of the residues forming the pair are used. To represent the latter subspace, Cartesian atomic coordinates of the 8 main chain atoms (N, $C_\alpha$, C and O) of the two residues are mapped to 16 inter-atomic distances between the residues. This gives 24 variables in total, including 8 torsional angles and 16 distances. For the torsional angle variables, the 1-d kernel function for determining neighbors takes the form,

$$\mathrm{h}(\tau^x, \tau^y) = \begin{cases} 1, & if |\tau^x - \tau^y| < \delta, and \\ \left[\frac{|\tau^x - \tau^y| - 1.5*\delta}{\delta - 1.5*\delta}\right]^2, & if \ \ \delta \leq |\tau^x - \tau^y| < 1.5 * \delta, \ and \\ 0, & if \ \ |\tau^x - \tau^y| < 1.5 * \delta. \end{cases} \quad \text{(S4)}$$

The values of the cutoff parameter $\delta$ have been determined adaptively and separately for the two main chain residues. For a given probing point projected onto the subspace of the 4 torsional angle variables of one residue, the cutoff parameter starts from the value of 40° for the flanking torsional angles and of 20° for the central torsional angle. Then the values of $\delta$ were iteratively increased (multiplied by a factor of 1.5 in each iteration) until the accumulated effective number of neighbors in this subspace exceeds 0.15. The 1-d kernel functions for the inter-atomic distance variables are of the form

$$\mathrm{h}(d^x, d^y) = \begin{cases} 1, & if \ \ |d^x - d^y| < \frac{d^x}{d_0} d^x, and \\ \left[2 - \frac{|d^x - d^y| * d_0}{d^x * d^x}\right]^2, & if \frac{d^x}{d_0} d^x \leq |d^x - d^y| < 2\frac{d^x}{d_0} d^x, \text{ and} \\ 0, & if \ \ |d^x - d^y| < 2\frac{d^x}{d_0} d^x. \end{cases} \quad \text{(S5)}$$

We note that with the above kernel function, the soft cutoff for neighbor counting in the 1-d distance space is given by $\frac{d^x}{d_0} d^x$, which increases with the square of the probing distance $d^x$. The purpose of this choice is to automatically lower the resolution of the energy function at longer distances. The initial value of $d_0$ was 100 Å and has been iteratively decreased (divided by a factor of 1.2 in each iteration) until the accumulated number of neighbors in the 16-dimensional subspace of the distance variables exceeds 6.0. We note that to reduce the computational cost associated with the adaptive determination of the cutoff parameters $\delta$ in (S4) and $d_0$ in (S5), their values have been determined on the basis of the numbers of neighbors in $S^r$, not in $S^t$. Neighbor counting in $S^r$ can be made very efficient by taking the advantage of the fact that underlying $S^r$, the overall "24"-dimensional distributions can be decomposed as the product of mutually independent distributions in three different subspaces, which include the two subspaces of the local conformational variables of the tow residues and the one subspace of the relative geometries between the two residues. Thus neighbor counting can be carried out separately in each of the subspaces. The cutoffs adaptively determined on the basis of $S^r$ can also be suitable for $S^t$ as the marginal distributions in the subspaces of the local conformational variables are the same between the reference and the training distributions. As for the subspace of the distance variables, although the marginal distributions are no longer the same between the training and the reference sample sets, the two marginal distributions contain the same intrinsic factor contributed by the non-isometric mapping from the Cartesian coordinates to the inter-atomic distances. Thus the values of $\delta$ and $d_0$ adaptively determined from $S^r$ have been used to carry out neighbor counting in $S^t$, during which all the 24 variables have been considered jointly (see formulae (10) and (11) in the main text).

The $e^{mc}_{non-local}$ estimated by NC were used to train the neural network $NN^{mc}_{non-local}$. Each of the local conformational variables (i.e., the backbone torsional angles) has been encoded with 6 trigonometric functions (formula (S1), with $k =$

1,2 and 4). Each of the 16 interatomic distances are encoded using the following 1-d Gaussian functions of varied centers,

$$d \rightarrow (e^{-\frac{(d-c_i)^2}{\sigma^2}}, i = 1,2,\ldots,8) \quad \text{(S6)}$$

The centers $c_i$ are at 2.5, 3.5, 4.5, 5.5, 6.5, 7.5, 9.0 and 10.5 Å, and the squared variance $\sigma^2$ is *0.5* Å$^2$. Besides the above input components, for each distance variable except for the $C_\alpha$-$C_\alpha$ distance, the difference between that distance and the $C_\alpha$-$C_\alpha$ distance has been included as an additional input component. These distance differences have been used directly as parts of the NN input because they encode the relative orientation between the two main chain residues straightforwardly. The total number of input nodes is $8 \times 6 + 16 \times 8 + 15$ =191. The training data for $NN^{mc}_{non-local}$ comprises 95000 points from $S^t$ and 195000 points from $S^r$ with corresponding NC-estimated $e^{mc}_{non-local}$ values. The chosen training points distribute evenly in the five $C_\alpha$-$C_\alpha$ distance ranges: 3 to 5.5 Å, 5.5 to 6.5 Å, 6.5 to 7.5 Å, 7.5 to 8.5 Å, and 8.5 to 9.5 Å. The resulting network has been tested on 5000 native points and 5000 computationally sampled points that have not been used in training the NN. The test points also distribute evenly in the different $C_\alpha$-$C_\alpha$ distance ranges. Using the RMS error on the test points as an indicator of the quality of the NN fitting, the number of nodes in the middle layer of the NN has been chosen to be 64.

**(4) The optional $e^{mc}_{local-HB}$ term**

As the local backbone conformations have been described in SCUBA by the Ramachandran angles and their combinations, the resulting energy is not sensitive to variances of local interatomic distances, such as the main chain local hydrogen bond distances. This leads some of the hydrogen bonds in the α helices to have wider distance distributions in the SD simulations as compared with the native structures (see Figure S9d). To remedy this, an optional $e_{local-HB}^{mc}$ term has been developed. This term depends on the configuration of 6 atoms from two dipeptide unit, $C_i, O_i, N_{i+1}, C_j, O_j,$ and $N_{j+1}$, in which i and j are the sequential residue numbers and $\mathrm{j} - \mathrm{i} \leq 5$. Based on which one of the two distances $\mathrm{r}_{\mathrm{N}_{\mathrm{i+1}} O_j}$ and $\mathrm{r}_{\mathrm{N}_{\mathrm{j+1}} O_i}$ is smaller, three atoms in one of the dipeptide unit are considered as the potential hydrogen bond donor group and respectively denoted as $C_d, O_d,$ and $N_d$. The remaining three atoms in the other dipeptide unit belong to the potential hydrogen bond acceptor group and are respectively denoted as $C_a, O_a$ and $N_a$. On the basis of the criterion $\mathrm{r}_{\mathrm{N}_\mathrm{d} O_a} < r_{N_a O_d}$, potential donor and acceptor groups can be unambiguously assigned. For the estimation of $e_{local-HB}^{mc}$ by NC, the training set $S^t$ consisted of configurations of main chain atom groups extracted from the training native proteins. Configurations in the reference set $S^r$ have been computationally generated by changing the coordinates of the donor group atoms (as a whole) with uniformly distributed random translations and rotations, the resulting configurations leading to steric clashes or too long $r_{N_d O_a}$ (>5.5 Å) excluded. The number of neighbors of a probing configuration have been counted using 6 geometrical variables including the Cartesian coordinates of $N_d$ in a local coordinate frame defined using $C_a, O_a$ and $N_a$ and the Cartesian coordinates of $O_a$ in a local coordinate frame defined using $C_d, O_d$ and $N_d$. For each variable, a 1-d kernel function of the same form as formula (S5) is employed, with the soft cutoff parameter increasing with the square of $r_{N_d O_a}$ and being adaptively changed.

To ensure that $e^{mc}_{local-HB}$ mainly reflects the contributions of local hydrogen bond interactions, two adjustments have been made in computing the final NC-estimated $e^{mc}_{local-HB}$. One adjustment leads the energy to be gradually switched off when $r_{N_dO_a} >$ 4 Å and strictly goes to zero after $r_{N_dO_a} \geq 4.5$ $\AA$. The other leads the computed energy not to exceed a certain maximum value. These adjustments have been implemented by using the following formula to compute the final NC-estimated $e^{mc}_{local-HB}$,

$$e^{mc}_{local-HB} = -ln\left[1 + \alpha(r_{N_dO_a})\left(\frac{N^t/N^r+\varepsilon}{1+\varepsilon} - 1\right)\right] \quad \text{(S7)}$$

in which the switching function $\alpha(r_{N_dO_a})$ takes the value of 1 for $r_{N_dO_a} < 4$ Å and changes continuously from 1 to 0 between $r_{N_dO_a} = 4$ and 4.5 Å. The value of $\varepsilon$ has been chosen to be 0.01, so that the maximum of $e^{mc}_{local-HB}$ cannot exceed $-ln\frac{0.01}{1.01}$.

The neural network model $NN^{mc}_{local-HB}$ has been trained using configurations in $S^t$ and $S^r$ with associated NC-estimated $e^{mc}_{local-HB}$ values. The input to $NN^{mc}_{local-HB}$ include the distance $r_{N_dO_a}$ encoded using a series of Gaussian functions (formula (S6) with the centers of Gaussian functions at 2.5, 2.75, 3.0, 3.25, 3.5, 4.0, 4.5, 5.0, 5.5, 6.0 and 6.5 Å and the variances $\sigma^2$of 0.015 Å$^2$ for the first four Gaussians and 0.05 Å$^2$ for the remaining Gaussians), the angles $\theta_{C_dN_dO_a}$ and $\theta_{C_aO_aN_d}$ each encoded by 6 trigonometric functions (formula (S1)), and the torsional angles $\tau_{O_dC_dN_dO_a}$ and $\tau_{N_aC_aO_aN_d}$, encoded as

$$\tau_{O_dC_dN_dO_a} \rightarrow (sin\theta_{C_dN_dO_a} sink\tau_{O_dC_dN_dO_a}, sin\theta_{C_dN_dO_a} cosk\tau_{O_dC_dN_dO_a}, k = 1,2,4),$$

$$\tau_{N_aC_aO_aN_d} \rightarrow (sin\theta_{C_aO_aN_d} sink\tau_{N_aC_aO_aN_d}, sin\theta_{C_aO_aN_d} cosk\tau_{N_aC_aO_aN_d}, k = 1,2,4). \quad \text{(S8)}$$

The multiplication of the trigonometric functions of a torsional $\tau$ by the sin function of the bond angle $\theta$ resolves the issue associated with ill-defined $\tau$ for $\theta$ approaching 180°. The middle layer of $NN^{mc}_{local-HB}$ consists of 32 nodes. The $NN^{mc}_{local-HB}$ has been trained using more than 700000 points and tested using more than 7000 points not involved in training.

**(5) The $e^{SC}_{rotamer}$ terms**

For each of the natural residue types with a rotatable sidechain chemical bond, an $e^{SC}_{rotamer}$ term has been defined to be dependent on the sidechain torsional angles as well as the backbone Ramachandran torsional angles. In the NC step, the training point set $S^t$ comprises conformers of the concerned residue type extracted from the training native proteins. The reference point set $S^r$ has the same marginal probability distribution in the Ramachandran subspace as $S^t$, but has independently and uniformly distributed sidechain torsional angles. The 1-d kernel functions are the same as defined in formula (S3), with $\delta_{min} = 8^o$ and $\delta_{max} = 30^o$.

In the NN step, each torsional angle variable of $e^{SC}_{rotamer}$ is encoded using the 6 trigonometric functions given in formula (S1). Training data for $NN^{SC}_{rotamer}$ comprises points taken from the corresponding $S^t$ and $S^r$ sets combined with the NC-estimated $e^{SC}_{rotamer}$ values. Test data have been taken from the same sets but not used in the NN training.

**(6) The $e^{sc}_{packing}$ terms**

In SCUBA, the $e^{sc}_{packing}$ terms are used mainly to represent the excluded volume effects of sidechains. It also provides non-specific attractive interactions to counterbalance the thermal expansion effects in finite temperature SD simulations.

Thus these terms have been simply chosen to be dependent on individual inter-atomic distances, each comprising a Lennard-Jones form repulsive region and an inverted Gaussian attractive region. Replacing the Lennard-Jones form attractive region by an inverted Gaussian form leads to a flatter bottom of the attractive well and much more rapid decay of the interaction at longer distances. The exact definition is

$$\mathrm{e}_{packing}^{\mathrm{sc}}(r)=\begin{cases}0.56*\left(\frac{\sigma^{12}}{r^{12}}-\frac{\sigma^{6}}{r^{6}}\right)+0.14-e_{min}, & if \;\; r<r_{min}, \;\; and\\ -e_{min}exp\left[-\left(\frac{r-r_{min}}{0.14*r_{min}}\right)^{2}\right], & if \;\; r\geq r_{min.}\end{cases} \tag{S9}$$

In formula (S9), $r_{min}$ is the location of the bottom of the well and $e_{min}$ is the depth of the well, and $\sigma=2^{-1/6}r_{min}$. The $r_{min}$ or the optimum packing distances are sums of atom type specific half packing distances (see Table S1 for their actual values). If both of the interacting atoms are non-polar, the value of $e_{min}$ is 1.0 scaled by a factor that depends on the total number of non-hydrogen atoms covalently bonded to the two interacting atoms (noted as $n_{conn}$), namely,

$$e_{min}=\epsilon*s(n_{conn}), \;\; with \;\; s(n_{conn})=\begin{cases}1.0 \;\; if \;\; n_{conn}\leq 2,\\ \frac{1.5}{n_{conn}-1} \;\; otherwise.\end{cases} \tag{S10}$$

The prefactor $\epsilon$ is 1.0 if both atoms are nonpolar. To roughly mimic the unfavorable desolvation of polar sidechain atoms, it is reduced to 0.2 when one of the two atoms is polar and to 0 when both atoms are polar. For hydrogen bond donor and acceptor pairs, $\epsilon$ is also 0 and $r_{min}$ is set to 2.9 Å. We note that SCUBA-driven backbone design is expected to be carried out with the simplified amino acid sequences, in which all sidechain atoms are non-polar.

We note that in the total SCUBA energy, $e_{packing}^{sc}$ is further scaled by the weight $w_{packing}^{sc}$ chosen based on SD simulations of native test proteins. Thus the above values of $\epsilon$ are only of relative meaning. We also note that parameters such as $r_{min}$ and the extension of the inverted Gaussian attractive well $(0.14 * r_{min})$ have been borrowed and simplified from the inter-atomic packing component of the ABACUS2 model, a model we developed for sequence design under given backbone structures. There, these parameters have been estimated on the basis of inter-atomic contact distance distributions in native proteins and refined by minimizing the differences between native and repacked sidechain conformations with fixed backbones.

**(7) The other energy terms**

The covalent terms $E^{mc}_{covalent}$ and $E^{sc}_{covalent}$ consists of harmonic bond length, bond angle, and improper dihedral angle terms as in usual molecular mechanics force fields, only that the equilibrium geometries and force constants have been estimated according to the distributions of corresponding geometric parameters in native proteins.

The steric term $E^{mc}_{steric}$ is a sum over main chain atom pairs separated by more than three covalent bonds. For each atom pair, the interaction is purely repulsive, taking the same form as the repulsive part of formula (S9). The atomic radius parameter σ takes the value of 3.1 Å for the $C_\alpha$ atoms, of 3.0 Å for the other atoms, and of 2.8 Å for hydrogen bonding atom pairs.

In some of the SD simulations reported in this work, user-defined restraining energy terms have been applied to heuristically restrict the conformational space accessible to SD sampling. These terms are described below.

i. The RMSD restraint

RMSD restraint has been applied to sample configurations around native structures that have been employed to calibrate the SCUBA energy weights (see below) . The restraining potential is a single one-sided harmonic function of the RMSD of the current configuration $\mathbf{r}$ with respect to the reference structure $\mathbf{r}_0$,

$$\mathrm{E}^{\mathrm{res}}_{RMSD}(\mathbf{r}) = \begin{cases} 0 & if \ \ RMSD(\mathbf{r}, \mathbf{r}_0) \le RMSD_0, \ and \\ \frac{1}{2} k^{res}_{\mathrm{RMSD}} [RMSD(\mathbf{r}, \mathbf{r}_0) - RMSD_0]^2, & \text{otherwise,} \end{cases} \tag{S11}$$

in which the parameter $RMSD_0$ defines the maximum structure deviation below which the configurations are unaffected by the restraint.

ii. The radius of gyration restraint

The radius of gyration ($R_g$) restraint has been introduced to bias the sampling to favor compact structures. It also serves to counteract the thermal expansion effects in the finite temperature SD simulations when the attractive interactions in the SCUBA energy terms are not strong enough for this purpose, e.g., when the simulation temperature is high or when a compact structure has not been formed in simulations starting from artificial backbones. To choose a functional form for the radius of gyration restraint, it is reasonable to assume that the effective volume available to a degree of freedom is proportional to the power of $R_g$ to a certain constant exponent. Then at a finite temperature, the free energy gain from volume expansion should be proportional to the logarithm of $R_g$. This leads to the choose of the following restraining potential on $R_g$ to compensate for the thermal expansion effects,

$$E^{res}_{R_g}(\boldsymbol{r}) = \begin{cases} 0, & \text{if } R_g(\boldsymbol{r}) \le R_g^0 \\ k^{res}_R ln \frac{R_{g(r)}}{R_g^0}, \text{otherwise.} \end{cases} \tag{S12}$$

in which the parameter $R_g^0$ defines the maximum $R_g$ below which the configurations are unaffected by the restraint. When $R_g^0$ is sufficiently small, its exact value becomes unimportant, because as long as $R_g(\mathbf{r})$ is larger than $R_g^0$, the restraining forces are not affected by the value of $R_g^0$. In addition, should a backbone structure becomes too compact, the logarithm dependence of $E_{R_g}^{res}(\boldsymbol{r})$ on $R_{g(\boldsymbol{r})}$ makes the forces derived from $E_{R_g}^{res}(\boldsymbol{r})$ to be very small in comparison with the steric repulsive forces. Thus we do not need to worry about that $E_{R_g}^{res}(\boldsymbol{r})$ could overly compress the structure even with very small $R_g^0$ and large $k_R^{res}$ (set to 300 Å$^{-2}$ in our calculations). In our SD simulations of the native test proteins at $T_r = 1.0$, $E_{R_g}^{res}$ have been applied with two choices of $R_g^0$ in two respective series of simulations. In one choice with $R_g^0$ =8 Å, a value that is significantly smaller than the $R_g$s of all the simulated structures, the restraint has been effectively turned on. In the other choice with $R_g^0$ being two times of the $R_g$ of the respective native structures, the restraint has been effectively turned off. In simulated annealing of the artificially constructed backbone structures, a small value of $R_g^0$ has been used to restrain the compactness of the sampled structures. In the latter simulations, $E_{R_g}^{res}(\boldsymbol{r})$ is needed to counterbalance the thermal expansion effects when the attractive sidechain packing is not yet considered, when well-packed structures is not yet formed, or when the simulation temperature is high.

iii. The secondary structure-specific local conformation restraint

In the initial stage SD simulations of the artificially constructed backbones (see below), the local conformations of backbone segments that are intended to form regular secondary structures in the final designed backbones have been restrained. The purpose was to restrict the conformational space to be searched by avoiding the disruption of the local conformations by poor initial structures and/or thermal fluctuations. For backbone segments desired to form helices, the Ramachandran torsional angles are restrained. For a segment desired to participate in a β-sheet, its extended conformation has been retained by restraining the Cα-Cα distances between all interlaced pairs of residues $i$ and $i+3$ within the segment to be longer than 10 Å. These restraints are enabled with one-sided harmonic potentials with sufficiently large force constants.

In the second stage of optimizing the artificially constructed backbones by SD simulations (see below), the secondary structure states should be stabilized by emerging favorable through-space interactions. Thus no secondary structure restraints were applied to avoid artificially biasing the final backbone structures.

**1.2 Calibrating the energy weights by SD simulations**

The objective of energy weight calibration has been to stabilize the native conformational states relative to conformations further away from the native structures. In addition, we would like the overall interaction strength to be as small as possible, so long as the tested native protein structures can retain their stability at $T_r = 1$. The calibration has been carried out using an approach that is conceptually similar to force field refinements using thermodynamics cycles in conformational space or “contract divergence”. Briefly, to refine a trial set of energy function parameters by this approach, two set of simulations using the trial parameter set may be carried out on chosen test proteins. One set comprises unrestrained simulations in which the structures are allowed to drift away from the starting native structure if the unrefined energy function cannot retain the stability of the native structure. In the other set of simulations, the structures are restrained to be close to the corresponding native structures (here by the

*RMSD* restraining energy described above, with the *RMSD0* parameter in formula (S11) set to 2.5 Å). Then the various energy components are compared between configurations sampled in the two sets of simulations to determine how an energy weight needs to be adjusted relative to its trial value. For example, if an energy component is systematically lower (higher) in the RMSD-unrestrained simulations than in the restrained ones, its weight should be reduced (increased) in the next round of test simulations. Otherwise the weight can be kept the same or tentatively reduced. The weights of the SCUBA energy components have been calibrated by this approach by first considering only two small globular test proteins, one all α (PDB ID 3l32) and another α+β (PDB ID 1a6j, chain A). The results have been further refined by considering 33 native proteins, which have been manually selected to be soluble globular ones of relatively small sizes, having relatively high-resolution structures determined by X-ray crystallography, containing no disulfide bond, and belonging to different fold classes. The PDB IDs of the native test proteins can be found in Figure 1 of the main text.

After we obtained a final set of energy weights ($w_{local}^{mc}$=2, $w_{non-local}^{mc} = 0.32$, $w_{rotamer}^{sc} = 2.4$, and $w_{packing}^{sc} = 3.1$), restraint-free SD simulations have been carried out with each of the individual weights systematically scaled by a factor of values between 0.2 and 2.0 with the other weights kept fixed. The results of these simulations (Figures S8 and S9) can indicate if further improvements of the model can be achieved by continuing to adjust only one weight. Besides these test simulations, simulations at $T_r = 1$ have been repeated with the total energy multiplied by an overall scaling factor

$S_{\text{total}}$ of values varied from 0.25 to 2.5. The results of these simulations (Figure S8d) can indicate whether the native structures are indeed located in minima on the SCUBA energy surface, and can suggest what may be the smallest overall interaction strength needed to maintain the stability of these minima against thermal fluctuations at $T_r = 1$.

In the above calibrating and testing SD simulations, the test proteins have been simulated with their native amino acid sequences or side chain types. To test whether the native backbones are stable under SCUBA without specific amino acid sequence information, additional SD simulations have been carried out on hypothetical proteins which retained the same starting backbone structures as the native proteins but are of non-native, simplified amino acid sequences, in which residues in α helices have been substituted by leucine, residues in β strands by valine, and residues in loops removed of sidechains (the secondary structure types have been assigned according to backbone-backbone hydrogen bond patterns in native structures).

If not specified otherwise, the SD integration time step is 2 *fs*, the atomic frictional coefficients are uniformly set to 0.5 $ps^{-1}$, the lengths of the test SD simulations on native proteins are 900 *ps*, the averaged RMSDs have been calculated on conformations sampled in the final 50 *ps* of the simulations, one snapshot per *ps* considered. If not specified, the RMSDs have been calculated on main chain atoms contained in regular secondary structure elements in the native structures.

#### 1.3 Optimizing the artificially constructed backbones by SD simulated annealing

The optimization process is divided into two stages. In the first stage, $w_{packing}^{sc}$ in the energy function has been set to 0, so that sidechain packing interactions have been excluded. Omitting sidechains leads to a smooth conformational energy surface so that an approximate minima of the main chain can be more easily reached. In addition, the secondary-structure-specific local conformation restraints (see above) have been applied to reduce the conformational search space at this stage. In the second stage, the secondary-structure-specific local conformation restraints have been removed. in addition, full sidechain interactions have been turned on, with the sidechain types determined according to the desired SS types of the backbone positions, namely, leucine for a helix position, valine for a strand position, and no sidechain for a loop position. In both stages, the optional local backbone hydrogen bond terms have been considered, and the radius of gyration restraint has been applied with a small $R_g^0$.

The first stage SD simulations mentioned above consist of a 10-ps relaxation simulation and a 60-ps simulated annealing simulation. In the relaxation simulation, a large frictional coefficient ($\gamma = 5\ ps^{-1}$) has been used with low temperature ($T_r = 0.1$) to eliminate strong unfavorable interactions in the artificially constructed initial structures, such as inter-atomic steric clashes (such interactions may lead to numerical instability in a simulation with smaller frictional coefficient). In the simulated annealing SD, the temperature has been changed between 2.0 and 0.5 in 6 cycles. Each cycle consists of a 4-*ps* period with $T_r = 2.0$, followed by a 1-*ps* transition period in which $T_r$ has been gradually changed from 2.0 to 0.5, and then by a 5-*ps* period in which $T_r$=0.5. The temperature course and lengths of the different periods have been determined on the basis of exploratory simulations to make sure that in the majority of the resulting conformations, the intended β-sheets are formed and the organizations of the SSEs are grossly consistent with the respective intended frameworks.

The second stage SD simulations consist of a 4-*ps* relaxation SD simulation with $\gamma = 5\ ps^{-1}$ and $T_r = 0.1$, followed by a 120-*ps* simulated annealing SD run with the temperature changed between 2.0 and 0.5 in 10 cycles. The resulting conformation has been further refined by another 120-*ps* simulated annealing SD run with the temperature cycled between 0.5 and 0.2. In each 10-*ps* simulated annealing cycle, the lengths of the high-temperature, the transition, and the low-temperature periods are 4 *ps*, 1 *ps*, and 5 *ps*, respectively.

When the above two-stage protocol is applied to an initial structure built according to a given intended framework containing β-sheet, the final structure may occasionally not contain the complete intended β-sheet. If this happened, we simply discarded that structure and tried the same protocol again on a new initial structure. For each intended framework, 10 final structures have been obtained. For different intended frameworks, the total number of initial structures that have been tried ranged between 10 to more than 30 to obtain the 10 final structures.

### 2. Supplementary Table

**Table S1.** Half optimum packing distances ($\frac{1}{2}r_{min}$) used to calculate $e^{sc}_{packing}$. For non-hydrogen-bonding atom pairs, the sum of the respective $\frac{1}{2}r_{min}$ values gives $r_{min}$. For hydrogen bonding atom pairs, the value of $r_{min}$ is 2.9 Å.

| Atom types | oxygen atoms | nitrogen atoms, bare carbon atoms; | $C_\alpha$ atoms, CH groups | $CH_2$ groups, sulfur atoms | $CH_3$ groups |
|---|---|---|---|---|---|
| $\frac{1}{2}r_{min}$(Å) | 1.6 | 1.7 | 1.85 | 1.9 | 1.93 |

### 3. Supplementary Figures

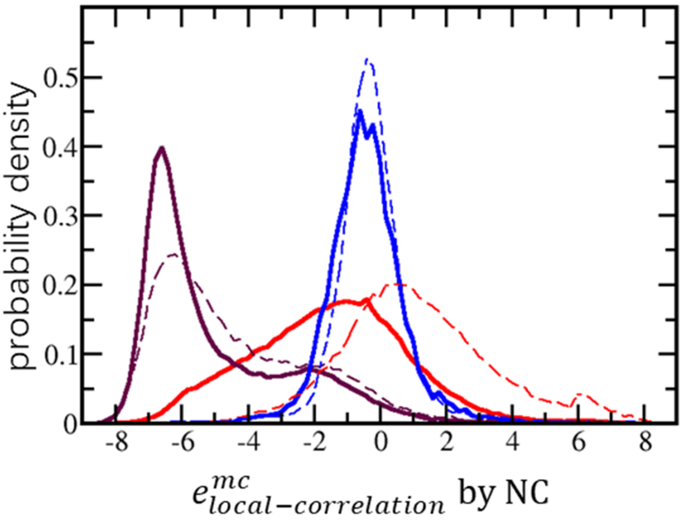


**Figure S1.** (a) The distributions of NC-estimated $e^{mc}_{local-correlation}$ of pentapeptide fragments in native protein structures (solid lines) and of randomly sampled pentapeptide conformations according to reference distributions in which the Ramachandran torsional angles of individual residues have been assumed to be independent from each other (dashed lines). Different colors correspond to pentapeptide fragments in different SS states. Purple: helix. Blue: strand, Red: coil. The distributions associated with random fragments are shifted to the higher energy side relative to the native fragments. The largest differences are for fragments in loops, for which the $e^{mc}_{local-correlation}$ showed broad distributions and large shifts from native to random, suggesting that to design loops with native-like backbones, it is important to consider the correlation between the backbone conformations of neighboring residues.

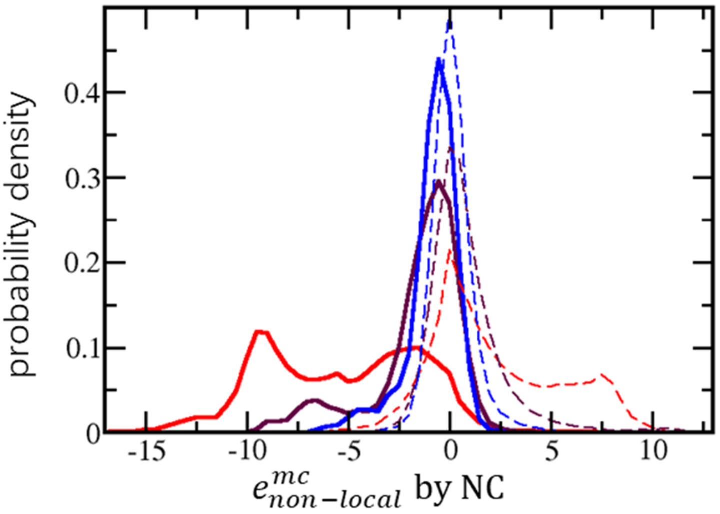


**Figure S2.** The distributions of NC-estimated $e^{mc}_{non-local}$ computed on main chain residue pair geometries in native proteins (solid lines) and sampled according to the reference distribution, in which the local Ramachandran torsional angles distribute independently between the residue pair, while the relative displacement and orientation between the residue pair follow random uniform distributions (dashed lines). Different colors correspond to different distance ranges. Red: $r_{C_\alpha - C_\alpha} < 6.5$ Å. Purple: 6.5 Å $< r_{C_\alpha - C_\alpha} \leq 8.5$ Å . Blue: 8.5 Å $< r_{C_\alpha - C_\alpha} \leq 9.5$ Å . While the energy distributions associated with random inter-residue geometries are, as expected, always shifted toward the higher energy side relative to the distributions computed on native structures, the shifts for distributions computed for geometries of shorter $r_{C_\alpha - C_\alpha}$ are larger and the corresponding distributions are wider, in consistence with the intuitive reasoning that the interaction strength increase with decreasing distance.

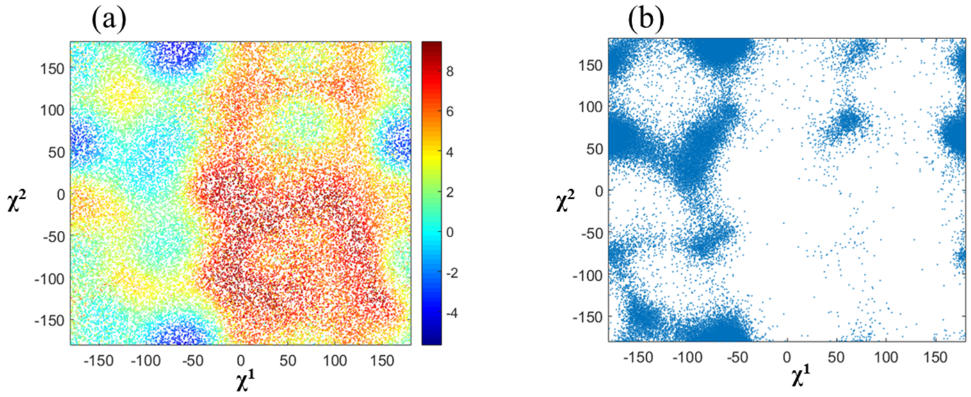


**Figure S3.** (a) The $e^{SC}_{rotamer}$ for leucine projected onto the plane spanned by the sidechain torsional angles $\chi^1$ and $\chi^2$. The points are the samples in $S^r$ and colored according to the NC-estimated energies. (b) Scattering plot of $\chi^1$ and $\chi^2$ of leucine residues in native proteins. The comparison between (a) and (b) indicates that the minima on $e^{SC}_{rotamer}$ captures peaks of the probability density faithfully. By definition, $e^{SC}_{rotamer}$ also depends on the backbone conformation. This causes in (a) the mixing of points of similar but not exactly the same colors within small areas on the $\chi^1$-$\chi^2$ plane.

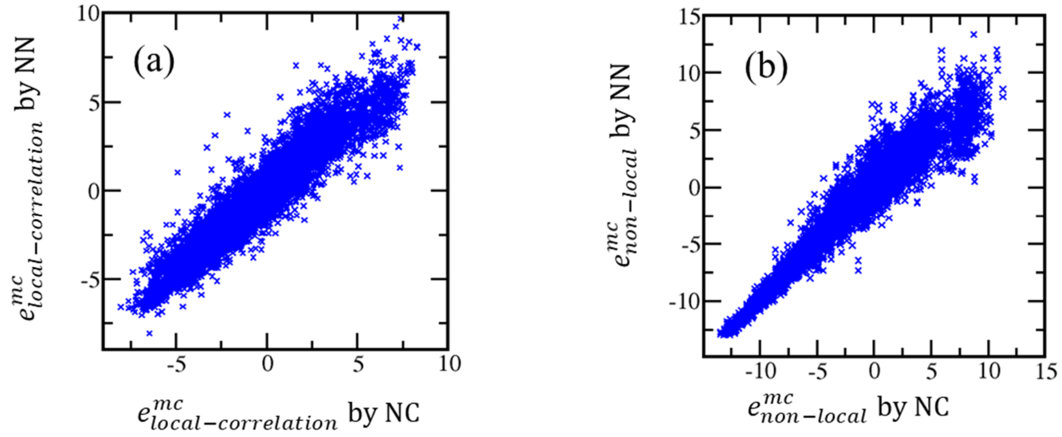


**Figure S4.** (a) NC-estimated values of $e^{mc}_{local-correlation}$ compared with the NN-estimated values. (b) NC-estimated values of $e^{mc}_{non-local}$ compared with the NN-estimated values. The geometric configurations at which the plotted energies have been evaluated have not been used to train the NN models. They consisted of approximately equal numbers of configurations extracted from native proteins and configurations computationally sampled according to respective reference distributions. The root mean square differences between the NC-estimated and the NN-predicted energy values are 0.7 and 0.8 (in reduced unit) for $e^{mc}_{local-correlation}$ and $e^{mc}_{non-local}$, respectively. Large portions of these errors should have come from the statistical noises which are contained in the NC-estimated energy values but may have been smoothened out in the analytical NN-predicted results. This interpretation about the error source is supported by that the magnitudes of the errors are largely unaffected by increasing the number of the NC-estimated points employed to train the NN model, or by tuning hyper-parameters of the NN model, including parameters defining the input-encoding function and the number of nodes in the middle layer.

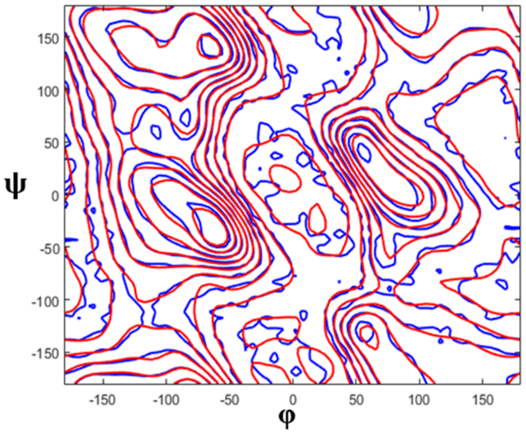


**Figure S5.** Contours of the NC-estimated (blue) and the NN-estimated （red） $e_{Rama}$.

The smoothening effect of the NN step can be directly visualized in this figure.

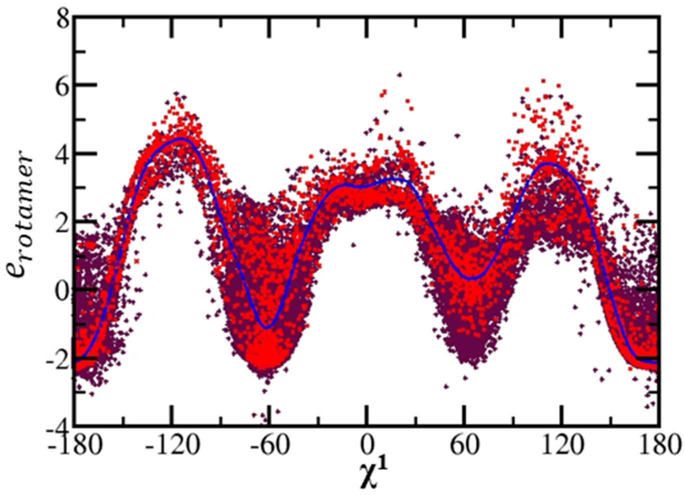


**Figure S6.** The $e^{SC}_{rotamer}$ of valine projected onto the $\chi^1$ axis. The points are from the set $S^t$. Purple dots and red dots are NC-estimated and NN-estimated values, respectively. The blue line is calculated using a NN model depending only on $\chi^1$ but not on the main chain torsional angles. Fitting the fluctuating NC-estimated $e^{SC}_{rotamer}$ of valine to only the $\chi^1$ variable by NN leads to a smooth one-dimensional energy curve. The actual NN-based $e^{SC}_{rotamer}$ values (red dots) depend additionally on the two main chain Ramachandran angles and do not follow a single-valued curve along $\chi^1$.

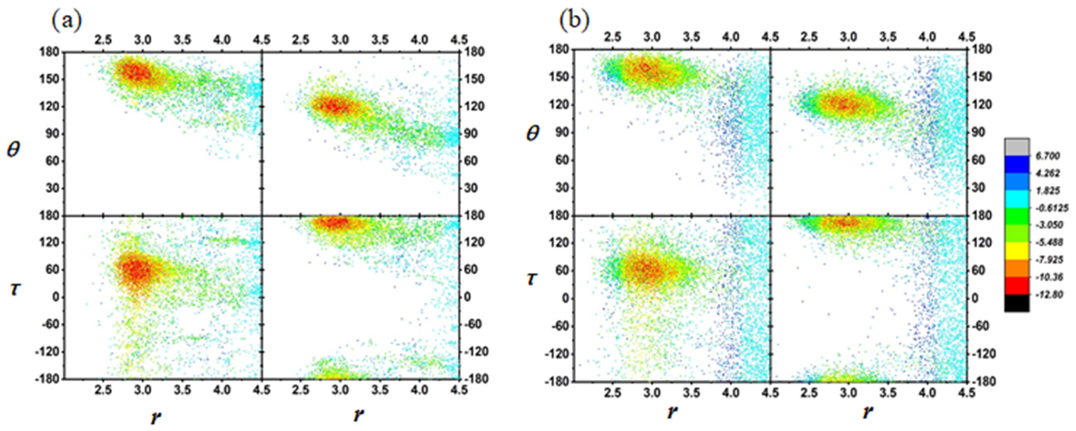


**Figure S7.** (a) The 2-d projections of native local main chain hydrogen bond geometries colored according to the NC-estimated $e_{local-HB}^{mc}$. The $r$ is the O-N distance. The $\theta$ is the C-O-N angle (top left) or C-N-O angle (top right). The $\tau$ is the C-O-N-C torsional angle (bottom left) or the C-N-O-C torsional angle (bottom right). (b) The same as (a) but for geometries computationally sampled according to a Boltzmann distribution depending on the NN-estimated $e_{local-HB}^{mc}$ and colored according to the same energy. The corresponding distributions and energy hyper-surfaces in (a) and (b) agree well, suggesting that the NC-NN statistical energy term can faithfully model a high-dimensional native distribution of a multiplex of strongly correlated geometric variables.

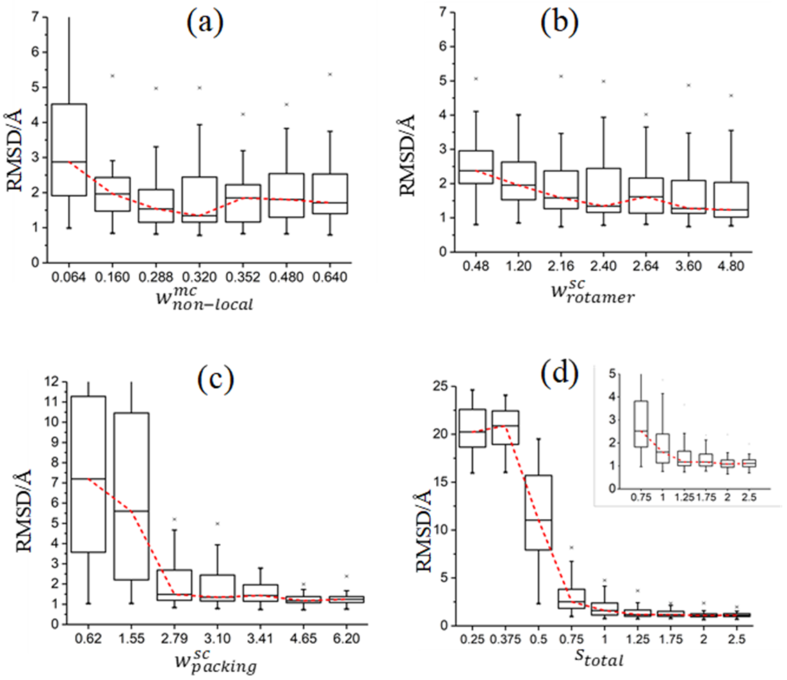


**Figure S8.** Box-and-whisker diagrams showing the medians, the interquartile ranges and the minimum and maximum deviations within 1.5 interquartile ranges of the averaged RMSDs from native structures in SD simulations of the 33 native test proteins. The trends of RMSD changing with varying individual weights (a to c) or a varying scaling factor of the total energy weights (d) are illustrated by the red dashed lines connecting the median values. The main chain local hydrogen bond terms have been turned on with $w^{mc}_{local-HB} = 0.6$. The radius of gyration restraint has been turned off. For $w^{mc}_{non-local}$, the chosen value of 0.32 is close to the optimum according to the RMSDs. With the increase of $w^{sc}_{rotamer}$ or $w^{sc}_{packing}$, the RMSDs first decrease and then flatten off. The final values of 2.4 for $w^{sc}_{rotamer}$ and 3.1 for $w^{sc}_{packing}$ are close to the minimum weights above which the RMSDs no longer decrease. The RMSDs first drops with the increase of $S_{total}$ and then flatten off approximate after $S_{total} = 1$, suggesting that the chosen values correspond to the weakest interaction strength needed to maintain the structural stability of the test native proteins against thermal fluctuation at $T_r = 1.0$. In addition, the RMSDs do not increase with the further increase of $S_{total}$, suggesting the native backbones are located in or near to SCUBA minima.

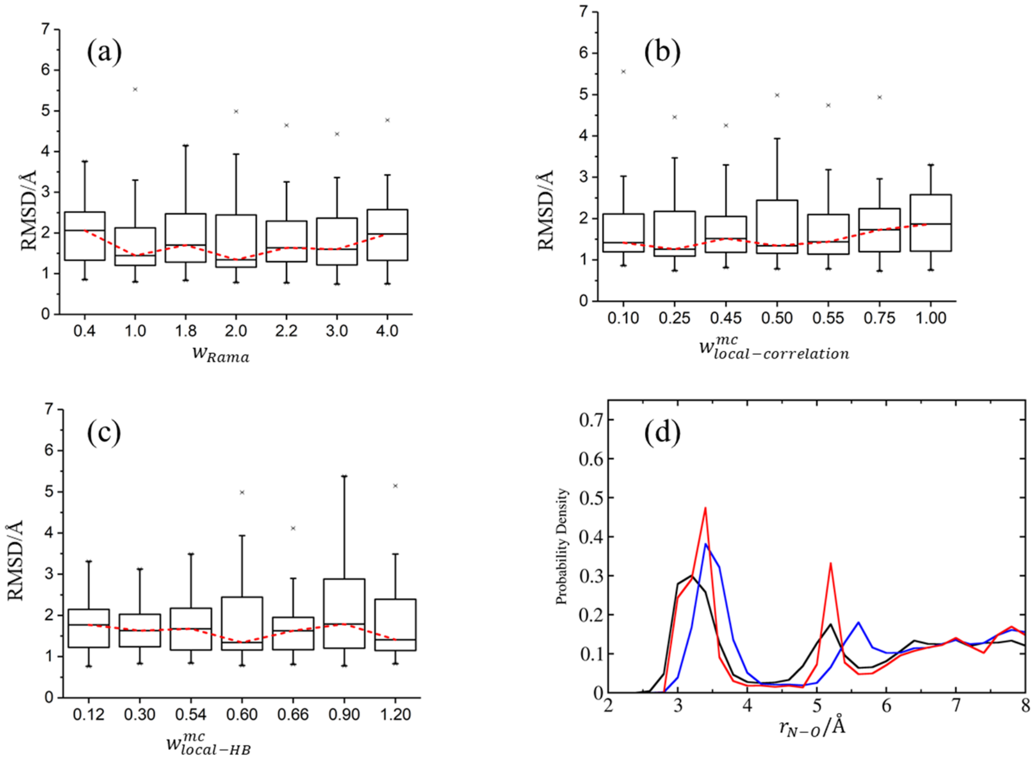


**Figure S9.** (a) to (c): box-and-whisker diagrams showing the medians, the interquartile ranges and the minimum and maximum deviations within 1.5 interquartile ranges of the averaged RMSDs from native structures in SD simulations of the 33 native test proteins. The trends of RMSD changing with varying different individual weights are shown, suggesting the monitored RMSDs are not sensitive to these weights. (d): The distribution of the N-O distance between residues *i* and residues *i+3, i+4* and *i+5* in the native structures (black), and of the averages of the same distances in SD simulations without (blue) and with (red) the optional main chain local hydrogen bond energy terms. Considering the main chain local hydrogen bond terms have notable effects on the distribution of the N-O distances. Without these energy terms, the peaks of the distribution of the distances averaged over the SD sampled configurations are shifted toward larger distances as compared with the native backbones. Turning on these terms can correct the locations of the peaks, although the peaks are sharper than those of the native distance distribution.

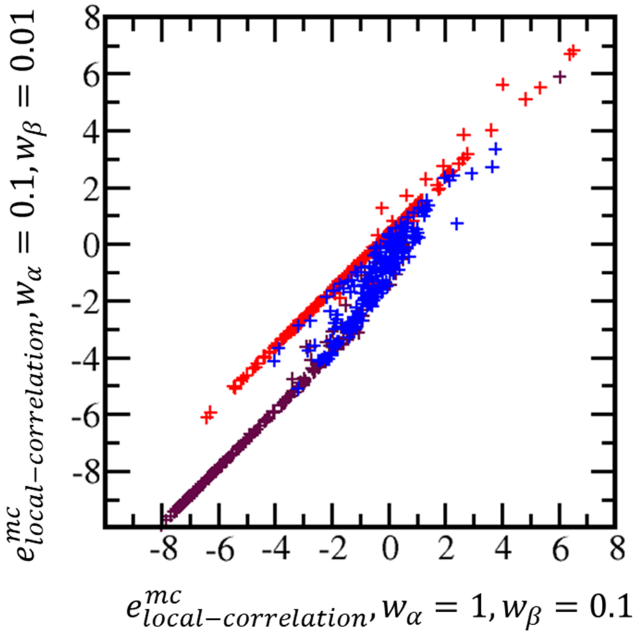


**Figure S10.** Comparisons of NC-estimated $e^{mc}_{local-correlation}$ of native fragments determined with different values for $w_\alpha$ and $w_\beta$. Different colors correspond to fragments in different local conformational states: brown for helix, blue for strand and red for coil.
